# Long-term memory formation depends on an astrocyte-to-neuron H_2_O_2_ signaling

**DOI:** 10.1101/2023.07.11.548505

**Authors:** Yasmine Rabah, Nisrine Sagar, Laure Pasquer, Pierre-Yves Plaçais, Thomas Preat

**Affiliations:** Energy & Memory, Brain Plasticity Unit, CNRS, ESPCI Paris, PSL Research University, Paris, France

## Abstract

Astrocytes interact with neurons during cognitive processes^1^. In particular, astrocytes help neurons to fight oxidative stress^2^, a needed function since active neurons are prone to reactive oxygen species (ROS) damage due to their high mitochondrial activity and relatively poor antioxidant defenses^3^. ROS also play major physiological functions^4,5^, but it remains unknown how neuronal ROS signaling is activated during memory formation and if astrocytes play a role in that process. We discovered in Drosophila an astrocyte-to-neuron H_2_O_2_ signaling cascade essential for long-term memory formation. Stimulation of astrocytes by acetylcholine induces an intracellular calcium increase that triggers the formation of extracellular O_2_°^−^ by astrocytic NADPH oxidase. Superoxide dismutase 3 secreted by astrocytes converts O_2_°^−^ into H_2_O_2_, which is imported into neurons of the olfactory memory center (the mushroom body), as revealed by *in vivo* H_2_O_2_ imaging using an ultrasensitive sensor. Importantly, SOD3 activity requires Cu^2+^, which we show is delivered by the neuronal Amyloid Precursor Protein. Furthermore, we found that human amyloid-ß peptide, involved in Alzheimer’s disease, inhibits the astrocytic cholinergic receptor and hampers memory formation by preventing H_2_O_2_ import into neurons. These findings could have major implications for the understanding of Alzheimer’s disease etiology, as soluble synaptic Aß42 correlates better with the pattern of cognitive decline in AD than amyloid plaques^6^, and since early pathology in cholinergic neurons of the basal forebrain predicts memory defects^7,8^.

## INTRODUCTION

Reactive oxygen species (ROS) such as superoxide (O_2_°^−^) and hydrogen peroxide (H_2_O_2_) are distinctive among metabolites because of their physiological ambivalence. On the one hand, ROS are toxic byproducts that can irreversibly oxidize macromolecules, and powerful scavenging enzymes are active to protect cells from ROS damage^9^. ROS production is linked to a great extent to mitochondrial activity, and neurons are especially prone to ROS damage due to their high mitochondrial activity and relatively poor antioxidant defense^10^. On the other hand, ROS play a positive role by activating signaling pathways involved in physiological functions such as cell survival, growth, or response to stress^5^. ROS signaling occurs mainly through reversible oxidation of redox sensitive cysteines, which requires a sharply regulated spatiotemporal increase in ROS concentration^11^. *In vitro* studies of long-term potentiation have shown that neuronal ROS signaling plays a positive role in neuronal synaptic plasticity^12,13^. Recent *in vivo* studies in Drosophila have shown that ROS signaling is required during development for activity-dependent plasticity in motoneurons^4,14^. While pioneering studies have reported that mutant mice with altered ROS metabolism display learning or memory defects^10,15^, it remains unknown how ROS signaling is recruited *in vivo* for memory formation. Astrocytes are known to regulate neuronal redox homeostasis^2^, but it is not known if they are involved in the initiation of neuronal ROS signaling for synaptic plasticity.

Building a mechanistic view of ROS signaling during memory formation is essential not only from a brain physiology perspective, but it could also improve our understanding of the onset of Alzheimer’s disease (AD), a neurodegenerative disease characterized in particular by memory loss and defects in redox homeostasis^16,17^. At the neuropathological level, AD is characterized by the progressive formation in the brain of neuritic plaques that correspond to the extracellular accumulation of amyloid beta (Aß) peptide, generated by cleavage of the transmembrane Amyloid Precursor Protein (APP), and by the formation of neurofibrillary tangles consisting of hyperphosphorylated TAU protein^18^. Early synaptic dysfunction has been associated with AD^17,19^, which correlates with cognitive decline^20^. However, the cascade of events leading to this fatal condition remains unclear.

Here, we have addressed these key brain physiology and pathology issues by combining brain H_2_O_2_ imaging using recently developed ultrasensitive sensors^21^ with functional characterization of ROS-related pathways during memory formation in Drosophila. We report that long-term memory (LTM) formation requires the establishment of a local H_2_O_2_ gradient in neurons of the olfactory memory center, which follows the activation of ROS-producing enzymes in astrocytes. This signaling cascade is impaired by AD-related Aß42. Our results provide a new framework to understand neuroglia interactions during memory formation in normal conditions and in AD.

## RESULTS

### Stimulated astrocytes generate extracellular ROS required for long-term memory

We initially observed that inhibiting NADPH oxidase (Nox) expression in adult astrocytes impairs olfactory LTM in Drosophila (Fig. 1A). The only function of the transmembrane enzyme Nox is to produce extracellular O_2_°^−^ from O_2_ and intracellular NADPH, and therefore our observation was potentially important from a ROS signaling perspective. For this experiment, we used a well-established paradigm of associative aversive olfactory conditioning^22,23^ named spaced conditioning (see Methods), along with inducible and spatially-controlled expression of a Nox-targeting RNAi. A single Nox exists in Drosophila, which shows strong sequence similarity with the human Nox5^24^. Like Nox5^11^, Drosophila Nox is activable by Ca^2+^ thanks to intracellular EF-hands domains that bind Ca^2+^ ^25^.

**Fig. 1.**
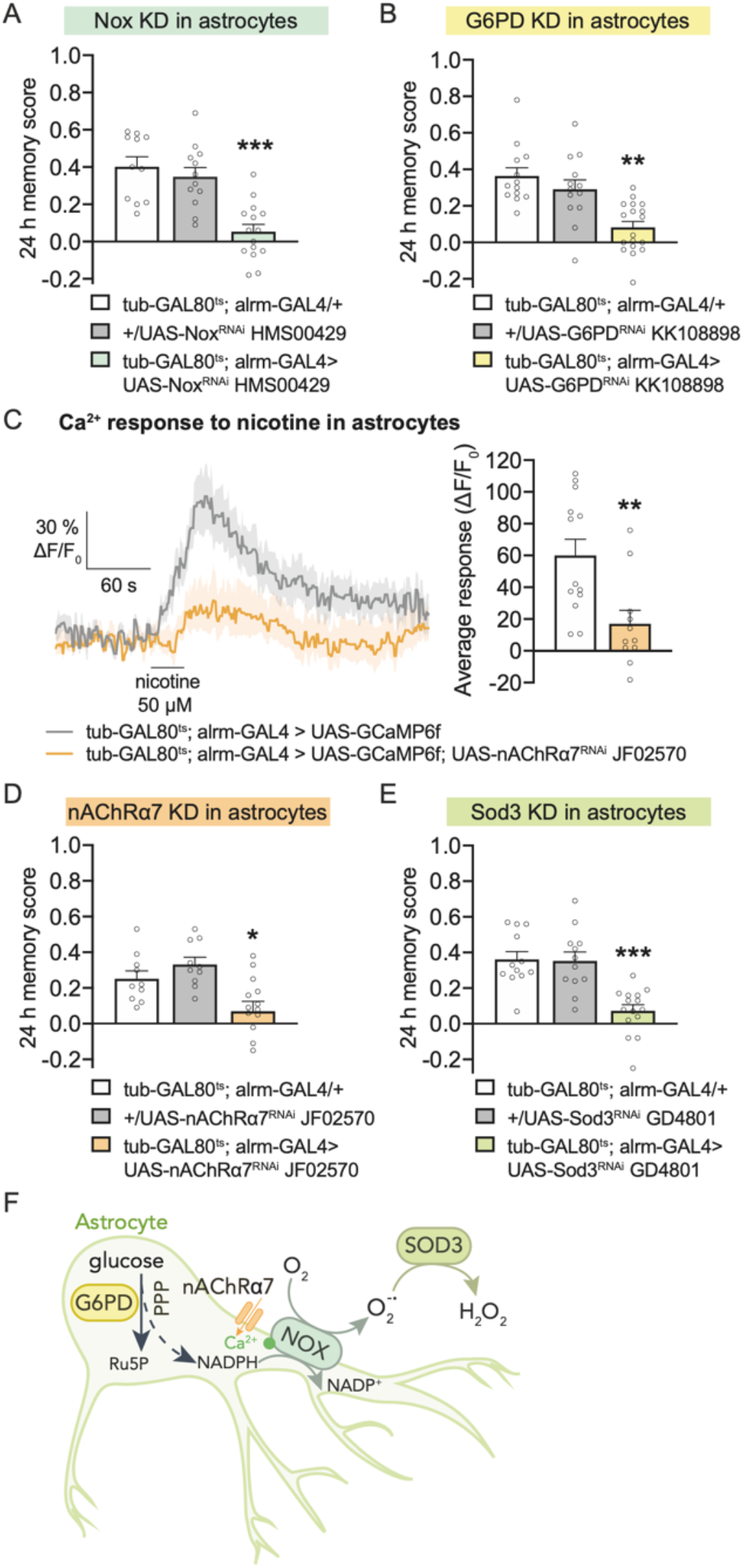
Astrocyte-derived H_2_O_2_ is required for LTM formation. (**A**) Nox knockdown (KD) in adult astrocytes impaired memory after 5x spaced conditioning (n = 11-15, F_2,35_ = 17.43, P < 0.001). (**B**) G6PD KD in adult astrocytes impaired memory after 5x spaced conditioning (n = 13-18, F_2,41_ = 12.98, P < 0.001). (**C**) The fluorescent calcium sensor GCaMP6f was expressed in adult astrocytes. Nicotine stimulation (50 µM) elicited calcium increase in astrocytes, which was significantly decreased in nAChRα7 KD (n = 11-13, t_22_ = 3.15, P = 0.005). (**D**) nAChRα7 KD in adult astrocytes impaired memory after 5x spaced conditioning (n = 10-14, F_2,31_ = 7.81, P = 0.002). (**E**) Sod3 KD in adult astrocytes impaired memory after 5x spaced conditioning (n = 12-15, F_2,36_ = 16.02, P < 0.001). (**F**) Scheme of the molecular actors involved in astrocytes for LTM. Nox, activated by calcium entry through nAChRa7, produces superoxide O_2_°^−^ from O_2_ using the NADPH co-factor derived from the pentose phosphate pathway (PPP). Extracellular O_2_°^−^ is converted to H_2_O_2_ by extracellular SOD3 secreted from the astrocytes. Ru5P: ribulose-5-phosphate. Data are presented as mean ± SEM. Significance level of t-test or least significant pairwise comparison following ANOVA: *P < 0.05; **P < 0.01; ***P < 0.001; ns: not significant, P > 0.05.

LTM formation requiring Nox activity in astrocytes depends on *de novo* protein synthesis^23^. Conversely, anesthesia-resistant memory (ARM), which forms after massed conditioning (see Methods) and does not depend on *de novo* protein synthesis, was not affected by Nox RNAi in adult astrocytes (Extended Data Fig. 1A). In addition, Nox RNAi did not affect perception of the stimuli used for conditioning (Extended Data Table 1), and LTM was normal when RNAi expression was not induced (Extended Data Fig. 1A). We confirmed the involvement of Nox in adult astrocytes for LTM with a second non-overlapping Nox RNAi (Extended Data Fig. 1B). These controls were performed systematically throughout this study, and for text simplification, RNAi knockdown displaying these phenotypes (impaired LTM with 2 RNAis, normal ARM, normal stimuli perception, normal LTM in absence of RNAi induction) will be simply reported as displaying a specific LTM defect.

The Nox substrate NADPH is mainly produced by the pentose phosphate pathway (PPP)^26^. To study this pathway we inhibited in adult astrocytes the expression of G6PD, the enzyme in the first step of the PPP^26^, and we observed a specific LTM defect (Fig. 1B). The essential role of astrocytic PPP in LTM formation was confirmed by inhibiting in astrocytes the expression of two other enzymes in this pathway, Pgd and Pgls^26^ (Extended Data Fig. 1, C-E, and Extended Data Table 1). Altogether, these results support the notion that Nox in astrocytes generates O_2_°^−^ required for LTM formation.

As stated above, similar to the human Nox5, Drosophila Nox is activated by Ca^2+^. Acetylcholine (Ach) is the major excitatory neurotransmitter of Drosophila brain neurons. Among the ionotropic cholinergic receptors, the channel formed of nAChRα7 subunits is the most permeable to calcium^27^, and is expressed in astrocytes in humans^28^. To examine whether nAChRα7 cholinergic stimulation induces a calcium flux in Drosophila astrocytes, we expressed the GCaMP6f calcium fluorescent reporter in astrocytes and stimulated the brains of live flies with nicotine, an nAChRα7 agonist. Nicotine stimulation induced an increased astrocytic calcium concentration in control flies (Fig. 1C). This effect was dampened when nAChRα7 expression was inhibited in astrocytes with RNAi (Fig. 1C). We next addressed the role of astrocytic nAChRα7 in memory and showed that nAchRα7 knockdown in adult astrocytes specifically impaired LTM (Fig. 1D and Extended Data Fig. 1F, G, and Extended Data Table 1). These results suggest that during spaced conditioning ACh release can activate calcium signaling in astrocytes and is required for LTM.

While O_2_°^−^ is a highly reactive and therefore toxic ROS with a short half-life, H_2_O_2_, which is less reactive and has a longer half-life, is more suitable for ROS signaling^9,29^. Nox activity in the plasma membrane produces extracellular O_2_°^−^, whose conversion into H_2_O_2_ is catalyzed by the only secreted superoxide dismutase (SOD), SOD3^30^. In the Drosophila brain, SOD3 is more strongly expressed in astrocytes than in neurons, as shown by single-cell transcriptomics data^31^ (Extended Data Fig. 2A). Inhibiting Sod3 expression in adult astrocytes with RNAi specifically impaired LTM (Fig. 1E and Extended Data Fig.2B, C, and Extended Data Table 1). These results suggest that a nAChRα7-NOX-SOD3 signaling cascade takes place upon LTM formation, which is triggered by Ach stimulation of astrocytes, resulting in extracellular H_2_O_2_ formation (Fig. 1F).

Astrocytic processes overlap with synapses in Drosophila brain^32^, as found in the tripartite synapse in the mammalian brain^33^. We therefore wondered if astrocytes could be a source of beneficial ROS imported by neurons during LTM formation. In particular, is H_2_O_2_ signaling involved during LTM formation in the mushroom body (MB) olfactory memory center?

### An H_2_O_2_ memory-related gradient forms in the mushroom body upon LTM formation

The MB is a bilateral structure consisting of about 2,000 cholinergic neurons^34^ in each brain hemisphere. Axons of the MB neurons form bundles and branch out, giving shape to anatomical structures called lobes that synapse with the MB output neurons. MB neurons are classified into three different subtypes: the α/β neurons, whose axons branch to form an α vertical projection and a medial β projection (Fig. 2A); the α’/β’ neurons; and the γ neurons, which form a single medial γ lobe. The axonal projection of the α lobe is specifically involved in LTM formation^22,35^. We aimed to image H_2_O_2_ in MB neurons upon LTM formation. Recently, the ability to observe physiological H_2_O_2_ levels has become possible with the development of an ultra-sensitive H_2_O_2_ fluorescent sensor, roGFP2-Tsa2ΔC_R_^21^, which we adapted for use in Drosophila (Fig. 2B and Extended Data Fig. 3A).

**Fig. 2.**
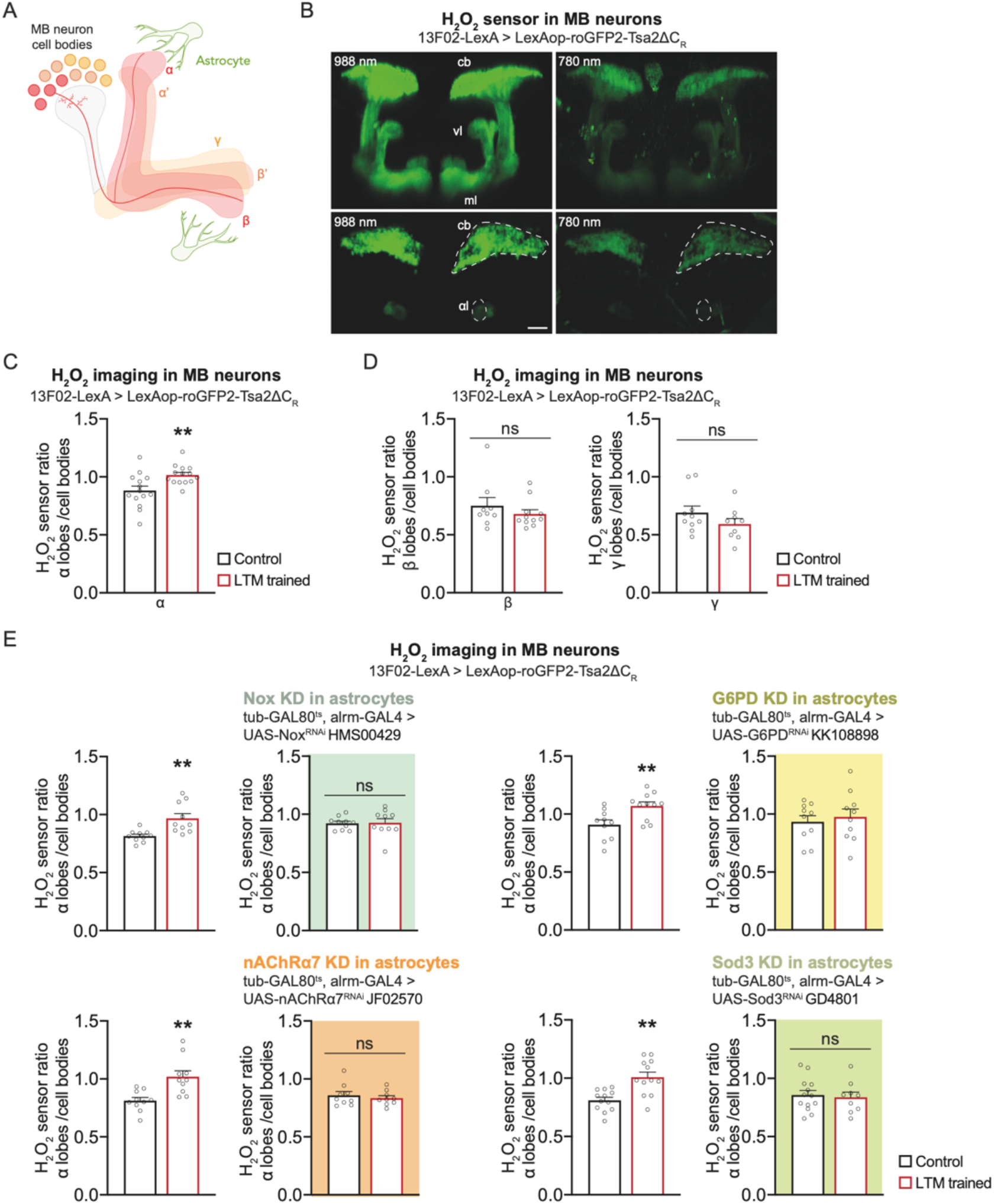
Astrocyte-derived H_2_O_2_ is imported in MB α lobes upon LTM formation. (**A**) Illustration of mushroom body structure. Axons of MB neurons follow a branching pattern that forms vertical lobes (α and α’) and medial lobes (β, β’ and γ). (**B**) Images of the roGFP2-Tsa2ΔC_R_ H_2_O_2_ sensor expressed in MB neurons. Top row: 3D reconstructions of image stacks (128 µm-thick stack, 1 µm Z-step) of the H_2_O_2_ sensor in MB neurons at the two excitation wavelengths (988 and 780 nm). Cb: cell bodies; vl: vertical lobes (α and α’); ml: medial lobes (β, β’ and γ). Bottom row: horizontal plane with regions of interest, cell bodies and α vertical lobes delimited with dashed lines. Scale bar: 30 µm. (**C**) The H_2_O_2_ level in α lobes normalized to the cell bodies value is increased upon spaced training, in comparison to the non-associative unpaired control (n = 13-14, t_25_ = 2.82, P = 0.009) (**D**) The H_2_O_2_ level in β or γ lobes normalized to the H_2_O_2_ level in cell bodies after spaced training was similar to the unpaired control (β: n = 9-11, t_18_ = 0.94, P = 0.36; γ: n = 9-10, t_17_ = 1.30, P = 0.21). (**E**) Measurements of H_2_O_2_ levels in MB neurons after spaced training in the context of astrocytic KDs. The increase in the α lobes/cell bodies H_2_O_2_ level elicited by spaced training was impaired in KD of Nox (n = 10, t_18_ = 0.11, P = 0.91; wild-type control: n = 10, t_18_ = 3.47, P = 0.003), G6PD (n = 10, t_18_ = 0.49, P = 0.63; wild-type control: n = 10-11, t_19_ = 3.01, P = 0.007), nAChRα7 (n = 9, t_16_ = 0.63, P = 0.54; wild-type control: n = 10, t_18_ = 3.57, P = 0.002) and Sod3 (n = 9-13, t_20_ = 0.33, P = 0.75; wild-type control: n = 12, t_22_ = 3.93, P < 0.001). Data are presented as mean ± SEM. Significance level of t-test or least significant pairwise comparison following ANOVA: *P < 0.05; **P < 0.01; ***P < 0.001; ns: not significant, P > 0.05.

To observe change in H_2_O_2_ dynamics, flies expressing the H_2_O_2_ sensor in the MB were imaged between 0.5 h and 2 h after LTM conditioning, a time window when early LTM-encoding events occur^36,37^. Control flies were submitted to non-associative unpaired treatment that does not allow memory formation. Interestingly, in flies trained for LTM, we observed an increased H_2_O_2_ level in the MB α lobe (Fig. 2C), and no increase in the MB β and γ medial lobes (Fig. 2D). An additional point must be considered in order to interpret the H_2_O_2_ imaging data. As we previously demonstrated, Drosophila LTM formation involves activation of the PPP in MB neurons, fueled by Glut1-mediated glucose import^36^. Since PPP activity generates the reducing agent NADPH, its activation in the MB is expected to result in a global decrease in ROS concentration (Extended Data Fig. 3B). Indeed, detailed examination of imaging data revealed a decreased H_2_O_2_ level in MB cell bodies in flies trained for LTM (Extended Data Fig. 3C). Interestingly, when glucose import was impaired in adult MB, the decreased H_2_O_2_ response in MB neuron cell bodies after spaced conditioning was abolished (Extended Data Fig. 3D). Altogether, these data reveal the existence in LTM-forming flies of an H_2_O_2_ gradient within the MB neurons, generated by H_2_O_2_ influx within the α vertical lobe. These results are in agreement with the fact that the α lobe is specifically involved in LTM formation.

To test whether astrocyte activity modulates H_2_O_2_ signaling in the MB LTM center, we imaged H_2_O_2_ in MB neurons while interfering with the H_2_O_2_-producing cascade in astrocytes. The increased level of H_2_O_2_ in the MB α lobe was lost by inhibiting the expression of either NOX, nAChRα7, G6PD or SOD3 in adult astrocytes (Fig. 2E). These results indicate that the H_2_O_2_ gradient formed in MB neurons after spaced conditioning originates from astrocytic NOX and SOD3 activity in response to ACh stimulation.

### A redox relay cascade is required in MB for LTM formation

How does H_2_O_2_ produced extracellularly by astrocytic enzymes enter MB neurons upon LTM formation? Although H_2_O_2_ can diffuse across the plasma membrane, H_2_O_2_ import is strongly facilitated by channels in the aquaporin protein family^38^. We tested the potential involvement of the main Drosophila aquaporin, AQP, in LTM (Fig. 3A). Inhibiting AQP expression by RNAi in adult MB α/β neurons specifically impaired LTM (Fig. 3B and Extended Data Fig. 4A, B, and Extended Data Table 2). The LTM defect following AQP expression inhibition was linked to a loss in the H_2_O_2_ gradient after conditioning (Fig. 3C).

**Fig. 3.**
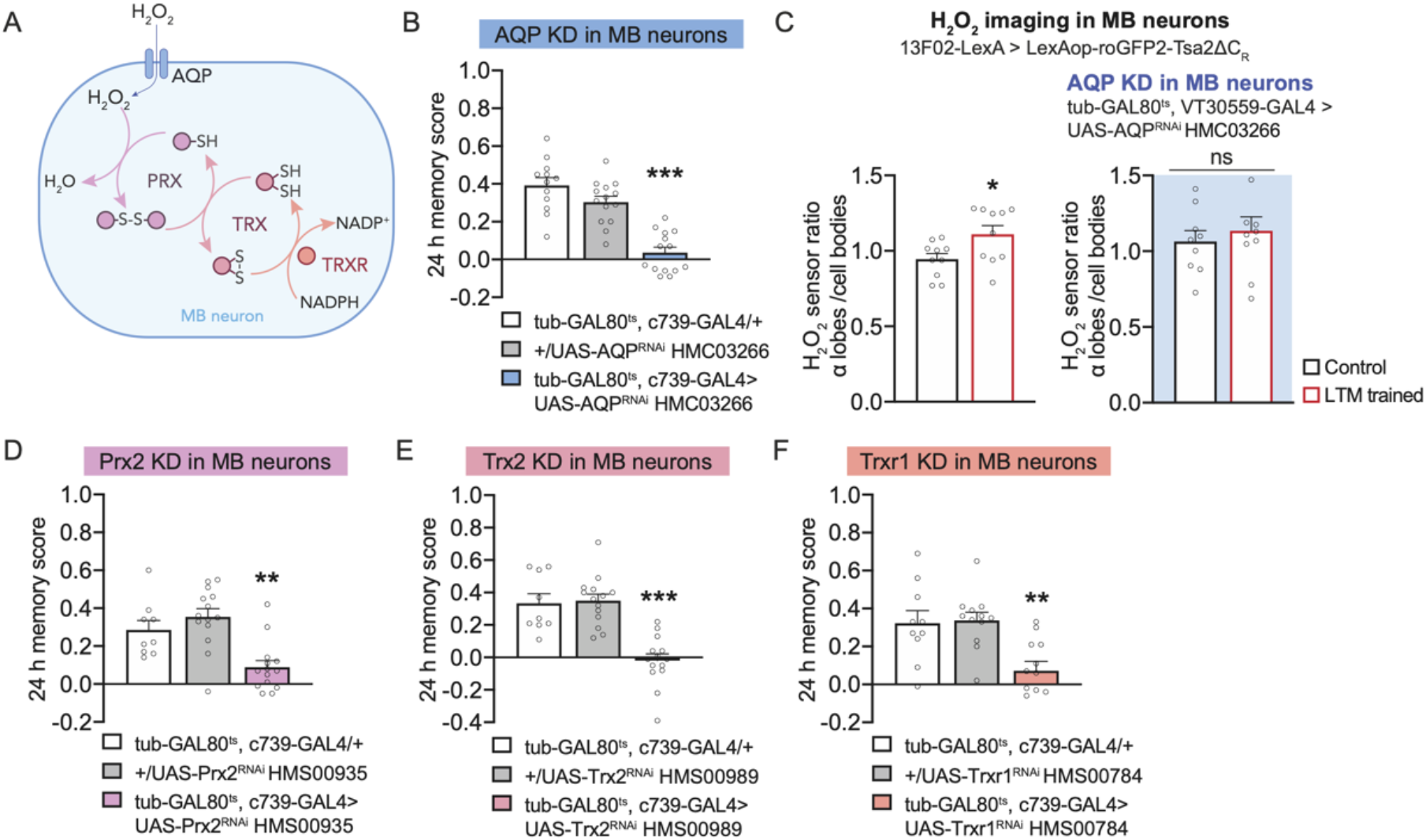
A redox-sensitive cascade is recruited upon H_2_O_2_ entry into MB neurons for LTM formation. (**A**) Putative redox relay cascade downstream of H_2_O_2_ import in MB neurons. (**B**) AQP KD in adult α/β MB neurons impaired memory after 5x spaced conditioning (n = 12-14, F_2,37_ = 31.24, P < 0.001). (**C**) The increase in H_2_O_2_ level in α lobes/cell bodies elicited by spaced training (n = 10, t_18_ = 2.42, P = 0.0261) was impaired in AQP KD in adult MB neurons (n = 9-10, t_17_ = 0.59, P = 0.56). (**D**) Prx2 KD in adult α/β MB neurons impaired memory after 5x spaced conditioning (n = 9-14, F_2,34_ = 12.08, P < 0.001). (**E**) Trx2 KD in adult α/β MB neurons impaired memory after 5x spaced conditioning (n = 12-14, F_2,37_ = 14.43, P < 0.001). (**F**) Trxr1 KD in adult α/β MB neurons impaired memory after 5x spaced conditioning (n = 10, F_2,27_ = 20.30, P < 0.001). Data are presented as mean ± SEM. Significance level of t-test or least significant pairwise comparison following ANOVA: *P < 0.05; **P < 0.01; ***P < 0.001; ns: not significant, P > 0.05.

These results support the idea that H_2_O_2_ signaling in α/β neurons occurs for LTM formation. H_2_O_2_ signaling can involve the direct oxidation of signaling proteins by H_2_O_2_. However, the physiological increase in local H_2_O_2_ concentration is generally considered to be too low to reach the required threshold, in particular because cytoplasmic catalase efficiently degrades H_2_O_239_. As an alternative mechanism, H_2_O_2_ signaling involves a cascade of redox-sensitive relay enzymes, starting with the Cys-based peroxidase Peroxiredoxin (Prx), an H_2_O_2_ scavenger (Fig. 3A). Oxidized Prx can interact with signaling proteins and modify their activity through reversible cysteine oxidation^40^. We identified Prx2 as the major Prx involved in LTM formation (Fig. 3D and Extended Data Fig. 5A, B, and Extended Data Table 3). Oxidized Prx are reduced by Thioredoxin (Trx)^40^, and we showed that Drosophila Trx2 is specifically involved in LTM (Fig. 3E and Extended Data Fig. 5C, D, and Extended Data Table 3). Lastly, Trx is reduced by a Trx reductase (Trxr) that oxidizes NADPH into NADP+ in the process. We identified Trxr1 as a major actor of LTM formation in α/ß neurons (Fig. 3F and Extended Data Fig. 5E, F, and Extended Data Table 3). Altogether, these results reveal a cascade of redox-sensitive enzymes required for LTM formation in α/β MB neurons. This cascade could be involved in the regulation of gene expression required for LTM formation^9,23^.

### Neuronal Amyloid Precursor Protein delivers extracellular copper for SOD3 activity during LTM formation

We have shown that H_2_O_2_ is generated extracellularly by SOD3, which requires extracellular Cu^2+^ for its catalytic activity^30^. Because of its potential toxicity, free Cu^2+^ is maintained at a very low concentration in cells and tissues by copper-binding proteins^41^. This raises the question of how sufficient Cu^2+^ levels can be accumulated at the MB synapses to sustain during LTM formation the activity of SOD3 secreted by astrocytes. We hypothesized that APPL, the single fly ortholog of Amyloid Precursor Protein (APP) transmembrane protein, might provide Cu^2+^ for astrocyte-derived SOD3 activity based on the following observations: i) the APP extracellular domain has several Cu^2+^ binding sites^42,43^ whose function remains poorly understood^44^, and Drosophila APPL carries in its conserved E2 ectodomain the 4 histidine that is characteristic of high-affinity Cu^2+^ binding sites^45^; ii) APP acts as an extracellular copper transporter in neurons, since APP overexpression decreases Cu^2+^ content in neurons *in vitro*, while APP loss-of-function leads to copper accumulation^46^; iii) APPL RNAi in MB neurons induces an LTM defect^47^; and iv) APPL exhibits neuronal expression and is more strongly expressed in α/β MB neurons^48^, which are involved in LTM^22,35^. We confirmed this strong α/β neuron expression with an HA-tagged Appl line generated by CRISPR (Fig. 4A and Extended Data Fig. 6A). Furthermore, we showed that knocking down APPL specifically in α/β neurons with RNAi induces an LTM defect (Fig. 4B and Extended Data Fig. 6B, C, and Extended Data Table 4).

**Fig. 4.**
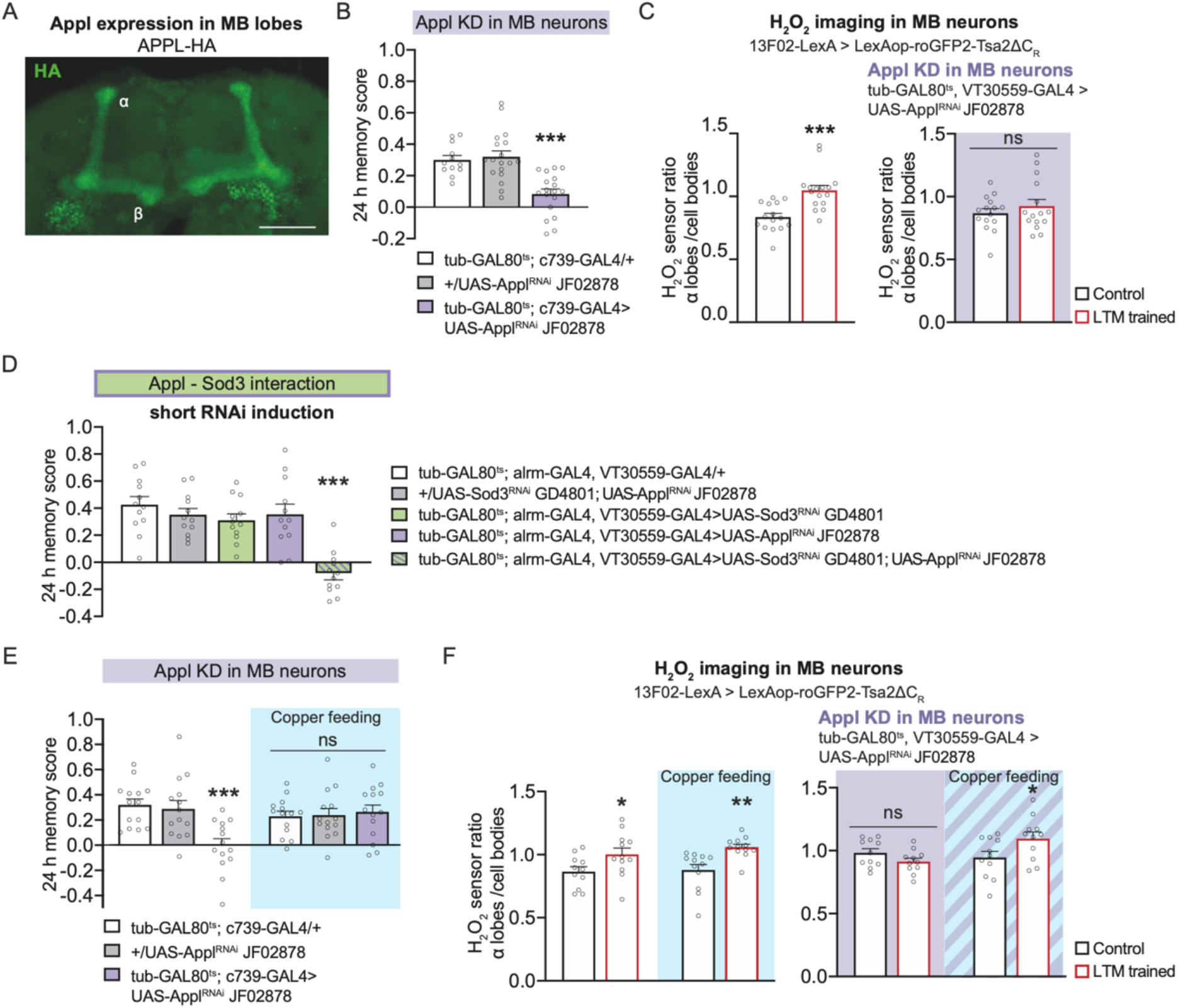
APPL provides copper to sustain SOD3 activity during LTM formation. **(A**) Image (maximum intensity projection of 22 µm-thick Z-stack acquisition, Z-step: 1 µm) of C-terminal HA-tagged APPL immunostaining showing the high expression of HA-tagged Appl in α/β MB neurons (green). Scale bar: 40 µm. (**B**) Appl KD in adult α/β neurons impaired memory after 5x spaced conditioning (n = 12-18, F_2,45_ = 15.58, P < 0.001). (**C**) The increase in H_2_O_2_ level in α lobes/cell bodies elicited by spaced training (n = 15-16, t_29_ = 4.30, P < 0.001) was impaired in Appl KD in adult MB neurons (n = 15, t_28_ = 0.88, P = 0.39). (**D**) Mild decreases in both Appl and Sod3 expression in MB neurons and astrocytes impaired memory after 5x spaced conditioning (n = 12, F_4,55_ = 13.22, P < 0.001). This mild decrease was obtained by activating RNAi expression for 1 day instead of 3 days. (**E**) Copper feeding (1 mM) 24 h before training restored normal memory capacity in flies with an Appl KD in adult α/β MB neurons (2-way ANOVA, n = 14, copper treatment: F_1,78_ = 0.94, P = 0.33; genotype: F_2,78_ = 4.48, P = 0.01; interaction: F_2,78_ = 6.70, P = 0.002). (**F**) Copper feeding (1 mM) 24 h before training restored the H_2_O_2_ level in α lobes/cell bodies in MB neuron presynaptic terminals of an Appl KD in adult MB neurons (right panel) (2-way ANOVA, n = 11-12, copper treatment: F_1,40_ = 3.07, P = 0.087; training: F_1,40_ = 0.92, P = 0.34; interaction: F_1,40_ = 7.01, P = 0.012). In wild-type controls (left panel), the increase in H_2_O_2_ level in the α lobes/cell bodies elicited by spaced training was not affected by copper feeding (2-way ANOVA, n = 11-12, copper treatment: F_1,43_ = 0.71, P = 0.40; training: F_1,43_ = 15.32, P < 0.001; interaction: F_1,43_ = 0.30, P = 0.59). Data are presented as mean ± SEM. Significance level of t-test or least significant pairwise comparison following ANOVA: *P < 0.05; **P < 0.01; ***P < 0.001; ns: not significant, P > 0.05.

If APPL indeed provides copper to SOD3, the H_2_O_2_ trace that forms upon spaced conditioning should require APPL. We therefore performed H_2_O_2_ *in vivo* imaging, and found that the H_2_O_2_ gradient was abolished when APPL expression was inhibited in adult MB (Fig. 4C). To further support the idea that APPL and SOD3 are involved in the same pathway, we searched for a synergistic interaction between Appl and Sod3 using mild inhibition of their expression. Inhibition of Appl or Sod3 expression for only 24 h in both MB neurons and astrocytes did not affect LTM formation (Fig. 4D). In contrast, LTM was specifically abolished when both Appl and Sod3 expressions were inhibited in MB neurons and astrocytes for 24 h (Fig. 4D and Extended Data Fig. 6D and Extended Data Table 4). Importantly, the simultaneous mild inhibition of Appl and Sod3 expressions in MB alone or astrocytes alone did not impact LTM (Extended Data Fig. 6E, F). Altogether, we can conclude that the inability to form LTM in double RNAi flies results from the simultaneous mild decrease of Appl expression in the MB, and of Sod3 expression in astrocytes. These results outline the existence of a strong functional interaction between APPL and SOD3, and are in agreement with the hypothesis that APPL supplies copper for SOD3 activity during LTM formation.

APP physiological function has been linked to many pathways^49,50^. To further demonstrate that the LTM defect of the APPL mutant is indeed due to a lack of copper, we aimed to rescue the LTM defect of the Appl mutant by feeding flies with copper-complemented food. Strikingly, the memory performance of the Appl mutant was fully rescued by 24-h, 1-mM copper feeding before conditioning, whereas the performance of the genotypic controls was unaffected (Fig. 4E). In agreement with this, the H_2_O_2_ gradient was restored after spaced conditioning in the Appl mutant supplemented with copper (Fig. 4F). Thus, copper-supplemented food can compensate for Appl knockdown at both the behavioral and H_2_O_2_ signaling levels. Altogether, these results strongly support the idea that the main function of APPL during LTM formation is to deliver extracellular copper for SOD3.

### Human Aß42 inhibits cholinergic receptor nAChRα7 in astrocytes and prevents H_2_O_2_ signaling during LTM formation

Following the observation that APPL is involved in the activation of H_2_O_2_ signaling, we wondered if, conversely, the toxic derivative of APP, amyloid beta (Aß), can inhibit this signaling pathway. Importantly, human Aß42 is a strong ligand for the nAChRα7 cholinergic receptor^51^. At very low concentrations (<100 pM), which we cannot generate in Drosophila using available genetic expression systems, oligomeric Aß42 was previously shown *in vitro* to activate nAChRα7^52^. However, in the 10-100 nM range, Aß42 becomes a strong inhibitor of nAChRα7, preventing activation of the receptor by its natural ligand, ACh^53^. Since we showed here that nAChRα7 activation in astrocytes initiates an H_2_O_2_ signaling pathway, we postulated that secreted Aß might impede this process. Because APPL shows no sequence similarity with APP at the level of the Aß sequence^54^, we overexpressed the human Aß42 peptide, as frequently performed in Drosophila to study the consequences of brain amyloid expression^55,56^. Thus, as previously reported, after several days of constitutive Aß42 expression, diffuse aggregates were observed in the Drosophila brain, along with neurodegenerative defects^55^. These defects are not observed after 3 days of Aß42 induction^55^. Since we aimed to evaluate the potential acute effect of Aß42 on astrocytic nAChRα7, expression was activated in MB α/ß neurons for a short 24-h period. We observed that the calcium response to nicotine was inhibited in the presence of Aß42 (Fig. 5A), suggesting that human Aß42 secreted by MB neurons inhibits Drosophila nAChRα7. We then showed that expressing Aß42 in MB for 24 h affected LTM specifically (Fig. 5B and Extended Data Fig. 7 and Extended Data Table 5). Lastly, we showed that Aß42 inhibits formation of the H_2_O_2_ gradient in MB neurons upon spaced conditioning (Fig. 5C). These results demonstrate that an acute expression of toxic Aß42 in young flies prevents H_2_O_2_ formation by astrocytes after spaced conditioning, and therefore LTM formation.

**Fig. 5.**
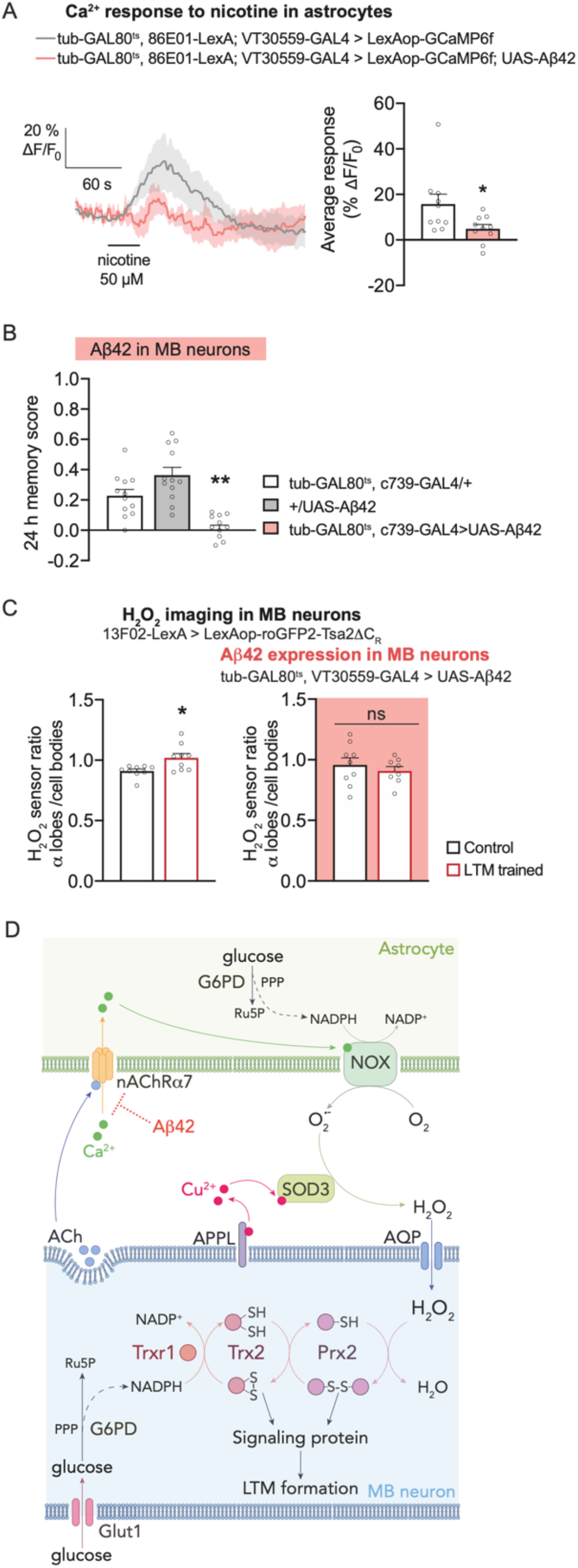
Human Aß42 inhibits astrocyte-neuron H_2_O_2_ signaling. (**A**) Nicotine stimulation (50 µM) elicited increased calcium in astrocytes, which was significantly decreased when human Aβ42 was expressed by MB neurons (n = 10, t_18_ = 2.29, P = 0.03). (**B**) Human Aβ42 expression in adult α/β neurons impaired memory after 5x spaced conditioning (n = 12, F_2,33_ = 18.51, P < 0.001). (**C**) Spaced training elicited an increase in the H_2_O_2_ level of the α lobes/cell bodies as compared to the unpaired control (n = 9, t_16_ = 2.75, P = 0.014), which was impaired by human Aβ42 expression by adult MB neurons (n = 8-9, t_15_ = 0.72, P = 0.48). **(D)** Model of <u>A</u>strocyte-to-<u>N</u>euron <u>H</u>_2_<u>O</u>_2_ <u>S</u>ignaling (ANHOS) for LTM formation. Acetylcholine release from MB neurons^34^ upon spaced training activates astrocytic nicotinic receptor nAChRα7, which induces a calcium elevation in astrocytes, resulting in NOX activation. NOX produces extracellular O_2_°^−^ in the presence of NADPH co-factor derived from astrocytic PPP. In the presence of Cu^2+^ delivered by MB neuronal APPL, extracellular astrocytic SOD3 converts O_2_°^−^ to H_2_O_2_. H_2_O_2_ enters MB neuron lobes through the AQP channel and fuels the redox-sensitive cascade composed of PRX2, TRX2 and TRXR1 enzymes. Regeneration of reduced forms of these enzymes is provided by PPP-derived NADPH produced during LTM formation^36^. Oxidized PRX2 and TRX2 can activate signaling cascades (to be identified), which allows LTM formation.

## DISCUSSION

ROS are byproducts produced mainly by mitochondria, and they are particularly toxic for neurons^10^. It has been suggested that ROS play a negative role in normal cognitive aging^57^, and that a defect in redox homeostasis is involved in the memory impairment characteristic of AD^57,58^. Glia help neurons fighting oxidative stress in various ways, including the production of reductive glutathione^2,59^ or accumulation of peroxidated lipids transferred from neurons^60^. However, ROS also play physiological roles^9^, and modulating the level of enzymes involved in ROS metabolism affects synaptic plasticity^10^ and learning and memory^15^. Nevertheless, the precise mechanisms controlling ROS signaling during memory formation have remained poorly known, in particular because imaging physiological ROS variations during memory formation was out of reach until the development of a genetically-encoded highly sensitive sensor^21^. Moreover, it was not known if glia could transfer beneficial ROS to neurons for neuronal plasticity and memory formation. Our data now provide a mechanistic understanding of the role of H_2_O_2_ signaling *in vivo* during memory formation. We have deciphered a pathway by which astrocytes deliver beneficial H_2_O_2_ to neurons for LTM formation, a phenomenon we have named Astrocyte-to-Neuron H_2_O_2_ Signaling (ANHOS) (Fig. 5D). The production of ROS by astrocytic NOX and SOD3 enzymes, which follows neurotransmitter-induced calcium increase, ensures a local H_2_O_2_ accumulation in the MB α lobe, the specific axonal projections encoding LTM^22,35^. Following H_2_O_2_ import, a signaling cascade is initiated by redox-sensitive enzymes^61^ comprising Prx2, Trx2 and TrxR1 (Fig. 5D). We propose that this cascade^5^ triggers LTM formation through reversible oxidation of specific cysteines in LTM-relevant signaling proteins. Although the future investigation of these molecular changes will be important, assessing transient changes of oxidation states is technically challenging, and this will require targeting the small subpopulation of α/β neurons that respond to the odorant used during LTM conditioning^62^.

SOD3 secreted by astrocytes plays a central role in ANHOS, as it generates the H_2_O_2_ molecules that will be imported by neurons. SOD3 requires a source of free copper for its catalytic activity. Although the free copper concentration remains extremely low in the brain because of its toxicity, copper concentrations around 100 µM have been reported at the synaptic cleft in mammals^63^. Our results in Drosophila suggest that an essential source of copper for the activity of SOD3 secreted by astrocytes comes from neuronal APPL, which exhibits an evolutionarily conserved copper-binding site on its extracellular domain. Indeed, APPL and SOD3 mutants display a strong synergistic interaction (Fig. 4D), and feeding copper to APPL flies rescued the LTM defect (Fig. 4E) and reestablished the H_2_O_2_ gradient in the olfactory memory center (Fig. 4F). Our results therefore suggest that the copper-binding property of APPL and its functional interaction with SOD3 are the major roles of APPL in LTM formation^47^. Interestingly, expression of extracellular SOD3 in the human brain is much stronger in astrocytes than in neurons^64^, as in Drosophila^31^ (Extended Data Fig. 2A). In mammals, APP binds copper^42,43^; it is expressed at the presynaptic active zone of neurons, where neurotransmitters are released^65,66^; and it is involved in synaptic plasticity and memory^50,67^. We propose that our model developed in Drosophila (Fig. 5D) can be generalized to this mammalian data, leading us to conclude that APP activates astrocytic SOD3 for memory formation in mammals.

From a pathological perspective, we show that human Aß42 blocks the ANHOS cascade by preventing Nox activation in astrocytes. We propose that this inhibitory process is at play during AD initiation. Indeed, several observations are in agreement with the pathophysiological hypothesis that impairment of astrocytes by diffusible Aß42 oligomers at cholinergic synapses may occur precociously in the AD brain: i) soluble synaptic Aß42 correlates better with the pattern of cognitive decline in AD than amyloid plaques^6^; ii) nAChRα7 is expressed by human brain astrocytes, in particular in the hippocampus^28^, a neural structure that plays a major role in episodic and spatial memory; iii) Aß42 inhibits nAChRα7 *in vitro* at a nM concentration^68^; and iv) in humans, pathology of the basal forebrain cholinergic neurons precedes cortical defects but is predictive of future memory deficits^7,8^. In addition, Sod3 has been associated with AD in several “omics” approaches, although its implication remains to be understood^69^. Our ANHOS model integrates observations of early AD and cholinergic neurons, and proposes a key role for SOD3 in AD. In normal conditions, cholinergic neurons of the basal forebrain, which send diffuse projections to many cortical areas as well as the hippocampus, are involved in memory formation^7,70^. The transition from a physiological condition to an AD condition might therefore involve a switch from an ANHOS activation by ACh to ANHOS inhibition via increased Aß42 production.

## Acknowledgments

We thank the Bloomington Drosophila Stock Center (NIH grant P40OD018537) and the Vienna Drosophila Resource Center for providing the flies used in this study. We thank Jean-Maurice Dura (Institute of Human Genetics, Montpellier, France) and Claude Desplan (New York University, USA) for fruitful discussions. We thank members of the Energy & Memory team for their comments on the manuscript.

## Author contributions

T.P. conceived and supervised the work; N.S. performed the behavioral analyses, with the support of Y.R. and L.P.; P.-Y.P. developed the assay for *in vivo* H_2_O_2_ imaging using the roGP2-Tsa2ΔC_R_ sensor; Y.R. performed H_2_O_2_ imaging and calcium imaging; L.P. characterized the Appl-HA line; N.S. performed the copper rescue experiment; T.P. wrote the main text of the manuscript; and Y.R. and P.-Y.P. contributed to writing the manuscript.

## Data and materials availability

All data are available in the manuscript or in the Extended Data section.

## Methods

### Experimental model

Flies (*Drosophila melanogaster*) were raised on standard medium at 18 or 23°C (depending on the experiments; see respective details below) and 60% humidity in a 12-h light/dark cycle. The study was performed on 0-3-day-old adult flies. For behavior experiments, both male and female flies were used. For imaging experiments, female flies were used because of their larger size. Transgenic flies were outcrossed for 5 generations to a reference strain carrying the w^1118^ mutation in an otherwise Canton-Special genetic background. Because TRiP RNAi transgenes are labelled by a y^+^ marker, these lines were outcrossed to a y^1^w^67c23^ strain in an otherwise Canton-Special background. All strains used in this study are described in Supplementary Table 1.

### Behavior Experiments

For behavior experiments, flies were raised on standard medium at 18°C and 60% humidity in a 12-h light/dark cycle. We used the TARGET system^71^ to inducibly express RNAi constructs exclusively in adult flies and not during development. To achieve the induction of RNAi expression, adult flies were kept at 30.5°C for 3 days before conditioning. To induce Aß42 expression (Aß42 carrying the pre-proenkephalin signal peptide for secretion^56^, adult flies were kept at 30.5°C for 24 h before conditioning. To test the interaction of Appl and Sod3, adult flies were kept at 30.5°C for 24 h only before conditioning, to achieve mild expression of RNAis. Otherwise, for non-induced experiments, experimental flies were transferred before conditioning to fresh bottles at 18°C.

All behavior experiments, including the sample sizes, were conducted similarly to other studies from our laboratory^22,36,37^. Groups of 20-50 flies were subjected to one of the following olfactory conditioning protocols at 25°C: five consecutive associative training cycles (5x massed training), or five associative cycles spaced by 15-min inter-trial intervals (5x spaced training). Non-associative control protocols (unpaired protocols) were also employed for imaging experiments. Conditioning was performed using previously described barrel-type machines that allow the parallel training of up to 6 groups. Throughout the conditioning protocol, each barrel was plugged into a constant air flow at 2 L.min^−1^. For a single cycle of associative training, flies were first exposed to an odorant (the CS+) for 1 min while 12 pulses of 5-s long 60 V electric shocks were delivered; flies were then exposed 45 s later to a second odorant without shocks (the CS-) for 1 min. The odorants 3-octanol (Fluka 74878, Sigma-Aldrich) and 4-methylcyclohexanol (Fluka 66360, Sigma-Aldrich), diluted in paraffin oil to a final concentration of 2.79·10^−1^ g.L^−1^, were alternately used as conditioned stimuli. During unpaired conditionings, the odor and shock stimuli were delivered separately in time, with shocks occurring 3 min before the first odorant.

Flies were kept at 18°C on standard medium between conditioning and the memory test. The memory test was performed in a T-maze apparatus^72^, 24 h after massed or spaced training, at 25°C. Each arm of the T-maze was connected to a bottle containing 3-octanol and 4-methylcyclohexanol, diluted in paraffin oil to a final concentration identical to the one used for conditioning. Flies were given 1 min to choose between either arm of the T-maze. A performance score was calculated as the number of flies avoiding the conditioned odor minus the number of flies preferring the conditioned odor, divided by the total number of flies. A single performance index value is the average of two scores obtained from two groups of genotypically identical flies conditioned in two reciprocal experiments, using either odorant (3-octanol or 4-methylcyclohexanol) as the CS^+^. The indicated ‘n’ is the number of independent performance index values for each genotype.

The shock response tests were performed at 25°C by placing flies in two connected compartments; electric shocks were provided in only one of the compartments. Flies were given 1 min to move freely in these compartments, after which they were trapped, collected, and counted. The compartment where the electric shocks were delivered was alternated between two consecutive groups. Shock avoidance was calculated as for the memory test.

Because the delivery of electric shocks can modify olfactory acuity, our olfactory avoidance tests were performed on flies that had first been presented another odor paired with electric shocks. Innate odor avoidance was measured in a T-maze similar to those used for memory tests, in which one arm of the T-maze was connected to a bottle with odor diluted in paraffin oil and the other arm was connected to a bottle with paraffin oil only. Naive flies were given the choice between the two arms during 1 min. The odor-interlaced side was alternated for successively tested groups. Odor concentrations used in this assay were the same as for the memory assays. At these concentrations, both odorants are innately repulsive.

### *In vivo* calcium imaging

Calcium imaging experiments were performed on flies expressing the GCaMP6f calcium sensor in astrocytes *via* the alrm-GAL4 driver, in combination with UAS-GCaMP6f. Transgenes were expressed in astrocytes using the inducible tub-GAL80^ts^; alrm-GAL4 driver. As done previously in our laboratory for imaging experiments^36,37^, flies were raised at 23°C to increase the expression level of genetically encoded sensors without allowing RNAi expression. Adult flies were kept at 30.5°C for 3 days before conditioning to achieve the induction of RNAi expression.

As in all previous imaging work from our laboratory, all *in vivo* imaging was performed on female flies, which are preferred since their larger size facilitates surgery. A single fly was picked and prepared for imaging as previously described^36,37^. Briefly, the head capsule was opened and the brain was exposed by gently removing the superior tracheae. The head capsule was bathed in artificial hemolymph solution for the duration of the preparation. The composition of this solution was: NaCl 130 mM (Sigma cat. # S9625), KCl 5 mM (Sigma cat. # P3911), MgCl_2_ 2 mM (Sigma cat. # M9272), CaCl_2_ 2 mM (Sigma cat. # C3881), D-trehalose 5 mM (Sigma cat. # 9531), sucrose 30 mM (Sigma cat. # S9378), and HEPES hemisodium salt 5 mM (Sigma cat. # H7637). At the end of surgery, any remaining solution was absorbed and a fresh 100-μL droplet of this solution was applied on top of the brain. Two-photon imaging was performed using a Leica TCS-SP5 upright microscope equipped with a 25x, 0.95 NA water-immersion objective. Two-photon excitation was achieved using a Mai Tai DeepSee laser tuned to 910 nm. The frame rate was 1 image per second. Recordings were acquired in the astrocytic region in the anterior part of the brain (where MB vertical lobes are located), at approximately 30 µm depth from the top of the brain.

Calcium imaging experiments with nicotine stimulation were performed as previously described^36^. Nicotine was freshly diluted from a commercial liquid (Sigma N3876) into the saline used for imaging on each experimental day. A perfusion setup at a flux of 2.5 mL·min^−1^ enabled the time-restricted application of 50 μM nicotine on top of the brain^36^. Baseline recording was performed during 1 min, after which the saline supply was switched to drug supply. The solution reached the *in vivo* preparation within 30 s. The stimulation was maintained for 30 s, before switching back to the saline perfusion for an additional 5 min. The GCaMP6f intensity was measured in the anterior part of the brain where MB vertical lobes are located. GCaMP6f signal was calculated over time after background subtraction and normalized by a baseline value calculated over the 30 s preceding drug injection using MATLAB software (MathWorks).

### *In vivo* H_2_O_2_ imaging

H_2_O_2_ imaging experiments were performed on flies expressing the excitation ratiometric H_2_O_2_ sensor roGFP2-Tsa2ΔC_R_^21^ in MB neurons *via* the 13F02-LexA driver, in combination with LexAop-roGFP2-Tsa2ΔC_R_ (generated in this study). Transgenes were expressed in astrocytes using the inducible tub-GAL80ts; alrm-GAL4 driver, or in MB neurons using the inducible tub-GAL80ts; VT30559-GAL4 driver. For imaging experiments, flies were raised at 23°C. Adult flies were kept at 30.5°C for 3 days before conditioning to achieve the induction of RNAi expression. For experiments on conditioned flies, data were collected indiscriminately from 30 min to 2 h after training.

Surgery was performed as for calcium imaging. At the end of surgery, any remaining solution was absorbed and a fresh 100-μL droplet of saline solution was applied on top of the brain. Two-photon imaging was performed using a pulsed infra-red laser (Insight X3, Spectra Physics), coupled to a Leica SP8 Dive upright microscope equipped with a 20x, 1.0 NA water-immersion objective and non-descanned spectral hybrid detectors. Emission was collected from 500 to 570 nm.

To characterize the roGFP2-Tsa2ΔC_R_ probe for 2-photon excitation microscopy, excitation spectra were measured from 780 to 1060 nm in steps of 8 nm. Oxidation of the probe was obtained by applying 2 mM H_2_O_2_ (final concentration) on top of the brain. Spectra in the basal and oxidized states were measured before and 15 min after H_2_O_2_ treatment, respectively. Excitation wavelengths of 780 nm and 988 nm were selected for further experiments, as they maximized the sensor emission ratio (ratio 780 nm/988 nm) between the basal and oxidized states. Experiments after conditioning were performed as follows: on a given brain area, two z-stacks (one at each acquisition wavelength) were consecutively acquired with a step of 1 µm and a line averaging of 3. The detector gain was strictly similar between the two stacks and was set to the 988 nm excitation wavelength at the highest value to avoid any pixel saturation. However, because of differential signal strength within different cellular compartments of MB neurons, two sets of stacks were acquired for each brain: one encompassing the MB lobes, and one covering the soma and calyx area, with different detector gains. For data analysis, the average intensity of 3 consecutive planes of the stack was calculated for each region of interest at the two excitation wavelengths. Oxidation of the sensor by H_2_O_2_ was measured by the ratio of the signal collected upon 780 nm excitation over the signal collected upon 988 nm excitation (referred to as “H_2_O_2_ sensor ratio”), using Leica Microsystems software LAS X Small (offline version).

### Dietary copper supplementation

Copper (II) sulfate pentahydrate (Thermo Fisher Scientific, cat. # 197730010) was added to melted standard food medium to achieve a final concentration of 1 mM, a concentration that does not affect survival rate under chronic exposure^73^. Flies were kept on regular food medium for 48 h at 30.5°C to achieve RNAi induction and then transferred to copper-enriched medium for 24 h at 30.5°C for both behavioral and H_2_O_2_ imaging experiments. No lethality was observed with this copper feeding diet. Control flies were transferred on regular medium for 24 h at 30.5°C.

### Immunohistochemistry

Before dissection, whole adult female flies (2-4 days old) were fixed in 4% paraformaldehyde in PBST (PBS containing 1% Triton X-100) at 4°C overnight. Brains were dissected in PBS solution and rinsed three times for 20 min in PBST, blocked with 2% bovine serum albumin (reference A9085, Sigma Aldrich) in PBST for 2 h at room temperature, and then incubated with primary antibodies at 1:200 (rat anti-HA, Roche) in the blocking buffer (2% BSA in PBST) at 4°C overnight. The following day, brains were rinsed 3 times for 20 min in PBST, and incubated with secondary antibodies at 1:400 (Invitrogen, goat anti-rat Alexa 488 Cat. A11006) in blocking buffer for 3 h at room temperature. Brains were further rinsed 3 times for 20 min in PBST, and were mounted in ProLong Mounting Medium (Lifetechnology) for imaging. Images were acquired with a Nikon A1R confocal microscope with a 20x objective.

### Western blot

Proteins were extracted from adult female heads after liquid nitrogen snap freezing and mechanical grinding in lysis buffer containing: sucrose 100 mM, KH_2_PO_4_/HPO_4_^−^ 40 mM, EDTA 30 mM, KCl 50 mM, 0.25% Triton X-100, DTT 10 mM, PMSF 0.5 mM, 1X protease inhibitors (Roche, cat. #11836153001). Protein extracts containing loading buffer NOVEX Tricine SDS sample buffer (Thermo Fisher Scientific, cat. # LC1676) were run in NOVEX 10-20% Tricine gels (Thermo Fisher, cat. # EC66252BOX) and then transferred on nitrocellulose membranes. Membranes were saturated with 15% non-fat milk, PBS-tween 0.2% for 1 h. Membranes were incubated with primary anti-HA antibodies diluted in PBS-tween 0.2% blocking medium (mouse anti-HA, 1:5000, Biolegend cat. # 901513) over night at 4°C under agitation. Membranes were rinsed 5 times for 8 min in PBS-tween 0.2% and incubated with secondary antibodies for 2 h at room temperature under agitation (HRP anti-mouse, 1:10000, Promega cat. # W4021). Membranes were further rinsed 5 times for 8 min in PBS-tween 0.2%. Revelation was conducted using NOVEX ECL substrate (Thermo Fisher, cat. WP20005) and chemiluminescence was detected with ImageQuant LAS4000. The same procedure was then applied to the same membranes with anti-tubulin antibodies (mouse anti-tubulin, 1:40000, Sigma Aldrich cat. # T16199). The molecular weight ladder corresponds to Invitrogen SeeBlue™ Plus2 (reference LC5925).

### Generation of transgenic flies

For the generation of the LexAop-roGFP2-Tsa2ΔC_R_ Drosophila line, the p415 TEF roGFP2-Tsa2ΔCR plasmid (Addgene #83238)^21^ was digested by NotI and XbaI. The resulting 1416 bp fragment was purified by electrophoresis and cloned into a pJFRC19 plasmid (13XLexAop2-IVS-myr::GFP addgene #26224)^74^. The resulting construct was verified by sequencing (the molecular cloning was outsourced to RD-Biotech, France). Transgenic fly strains were obtained by site-specific embryonic injection of the resulting vector in the attP18 landing site (chromosome 1), which was outsourced to Rainbow Transgenic Flies, Inc (CA, USA).

The Appl-HA line was generated using the CRISPR approach (outsourced to inDroso, FR). The 3xHA sequence (GCCGCCGTGTACCCCTACGACGTGCCCGACTACGCCGGCTACCCCTACGACGTGCCCGACTACGCCG GCTCCTACCCCTACGACGTGCCCGACTACGCCCCCGCCGCC) preceded by a linker (sequence: GGCGTGGGC) was inserted between the 11^th^ exon and the 3’UTR of the APPL gene using a guide RNA (5’ AAGTGAAA|GAGTAAGCGAGA3’). The genomic edition was strictly restricted to the 3xHA tag and linker, preventing any alterations due to the presence of a selection marker (scarless).

### Sod3 transcript levels

Relative expression of Sod3 in astrocytes and α/β neurons was obtained from published single-cell transcriptomic data^31^, according to clustering performed in the original study where astrocytes are found in cluster 10 and α/β in cluster 22.

### Quantification and statistical analysis

All data are presented as mean ± SEM. For behavior experiments, 2 groups of about 30 flies were reciprocally conditioned, using respectively octanol or methylcyclohexanol as the CS^+^. The memory score was calculated from the performance of two groups as described above, which represents a single experimental replicate. For imaging experiments, one replicate corresponds to one fly brain. Comparisons of the data series between two conditions were achieved by a two-tailed unpaired t-test. Comparisons between more than two distinct groups were made using a one-way ANOVA test, followed by Tukey pairwise comparisons between the experimental groups and their controls. ANOVA results are presented as the value of the Fisher distribution F(x,y) obtained from the data, where x is the number of degrees of freedom between groups and y is the total number of degrees of freedom for the distribution. For copper rescue experiments (Fig. 4, E and F), 2-way ANOVA tests were performed, followed by Sidak’s multiple comparisons test. Statistical tests were performed using the GraphPad Prism 8 software. In the figures, asterisks denote the significance level of the t-test, or of the least significant pairwise comparison following an ANOVA, with the following nomenclature: *P < 0.05; **P < 0.01; ***P < 0.001; ns: not significant, P > 0.05. Figures were designed using Adobe Illustrator.

