## Extended Data for "Long-term memory formation depends on an astrocyte-to-neuron H_2_O_2_ signaling"

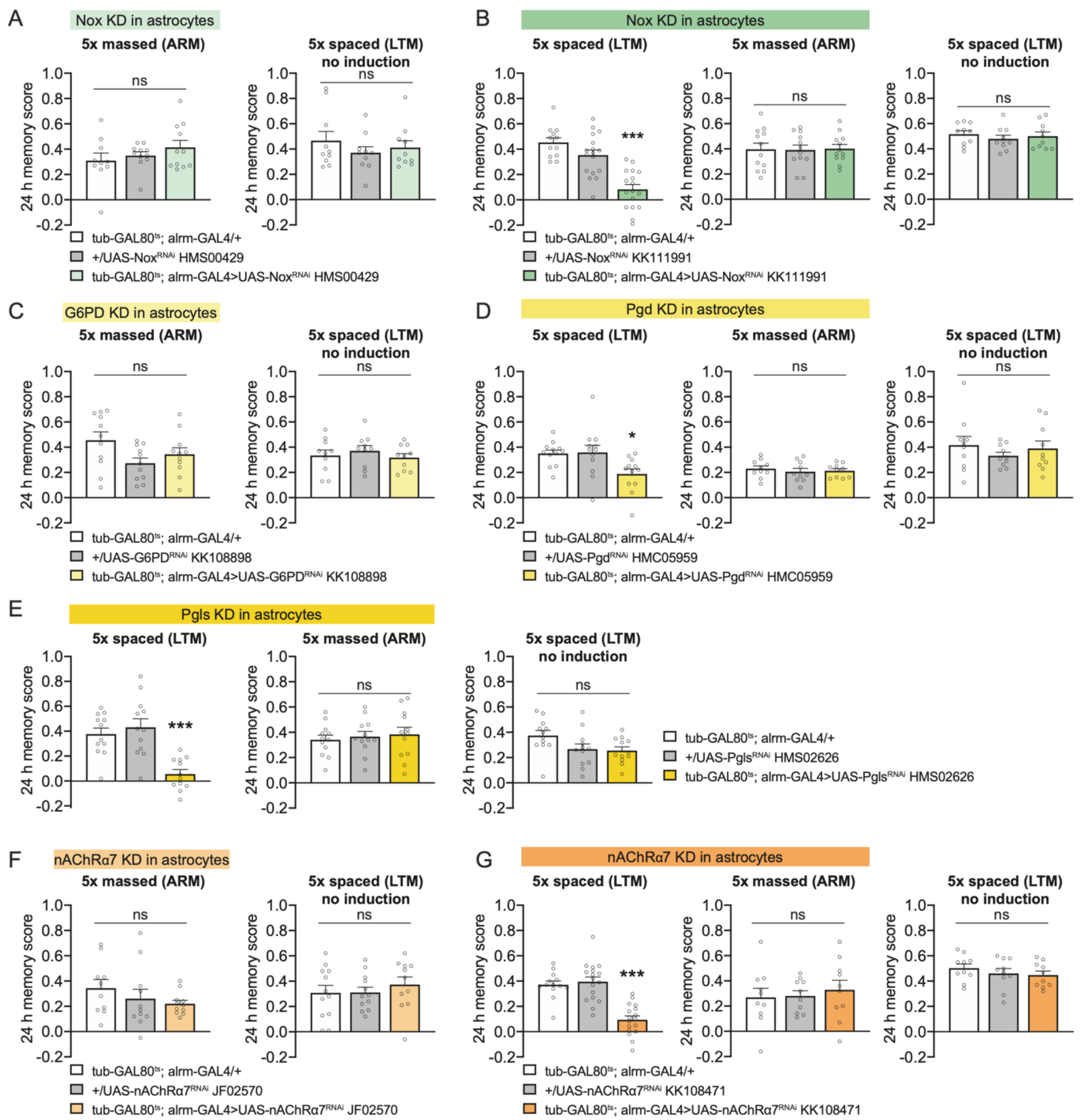


**Extended Data Fig. 1. Controls for behavioral experiments in Fig. 1**

**(A)** Memory after massed conditioning was not affected by Nox knockdown (KD) in adult astrocytes (n = 10-11, $F_{2,29}$ = 1.17, P = 0.32). Without Nox RNAi induction, memory after spaced training was normal (n = 10, $F_{2,27}$ = 0.67, P = 0.52). **(B)** The behavioral experiments were performed with a second RNAi against Nox. Nox KD in adult astrocytes impaired memory after spaced training (n = 12-17, $F_{2,43}$ = 24.03, P < 0.001), but did not affect memory after massed training (n = 12, $F_{2,33}$ = 0.018, P = 0.98). Without Nox RNAi induction, memory after spaced training was normal (n = 10, $F_{2,27}$ = 0.38, P = 0.68). **(C)** Memory after massed conditioning was not affected by G6PD KD in adult astrocytes (n = 11, $F_{2,30}$ = 2.93, P = 0.07). Without G6PD RNAi induction, memory after spaced training was normal (n = 10, $F_{2,27}$ = 0.49, P = 0.62). (**D)** Pgd KD in adult astrocytes impaired memory after spaced training (n = 12, $F_{2,33}$ = 4.92, P = 0.01), but did not affect memory after massed training (n = 10, $F_{2,27}$ = 0.31, P = 0.73). Without Pgd RNAi induction, memory after spaced training was normal (n = 10, $F_{2,27}$ = 0.61, P = 0.55). **(E)** Pgls KD in adult astrocytes impaired memory after spaced training (n = 12, $F_{2,33}$ = 14.75, P < 0.001), but did not affect memory after massed training (n = 12, $F_{2,33}$ = 0.24, P = 0.79). Without Pgls RNAi induction, memory after spaced training was normal (n = 12, $F_{2,33}$ = 3.11, P = 0.06). **(F)** Memory after massed conditioning was not affected by nAChRα7 KD in adult astrocytes (n = 10-11, $F_{2,29}$ = 1.07, P = 0.36). Without nAChRα7 RNAi induction, memory after spaced training was normal (n = 11-12, $F_{2,32}$ = 0.47, P = 0.63). (**G)** The behavioral experiments were performed with a second RNAi against nAChRα7. nAChRα7 KD in adult astrocytes impaired memory after spaced training (n = 13-17, $F_{2,44}$ = 26.95, P < 0.001), but did not affect memory after massed training (n = 10, $F_{2,27}$ = 0.24, P = 0.79). Without nAChRα7 RNAi induction, memory after spaced training was normal (n = 10, $F_{2,27}$ = 0.71, P = 0.50).

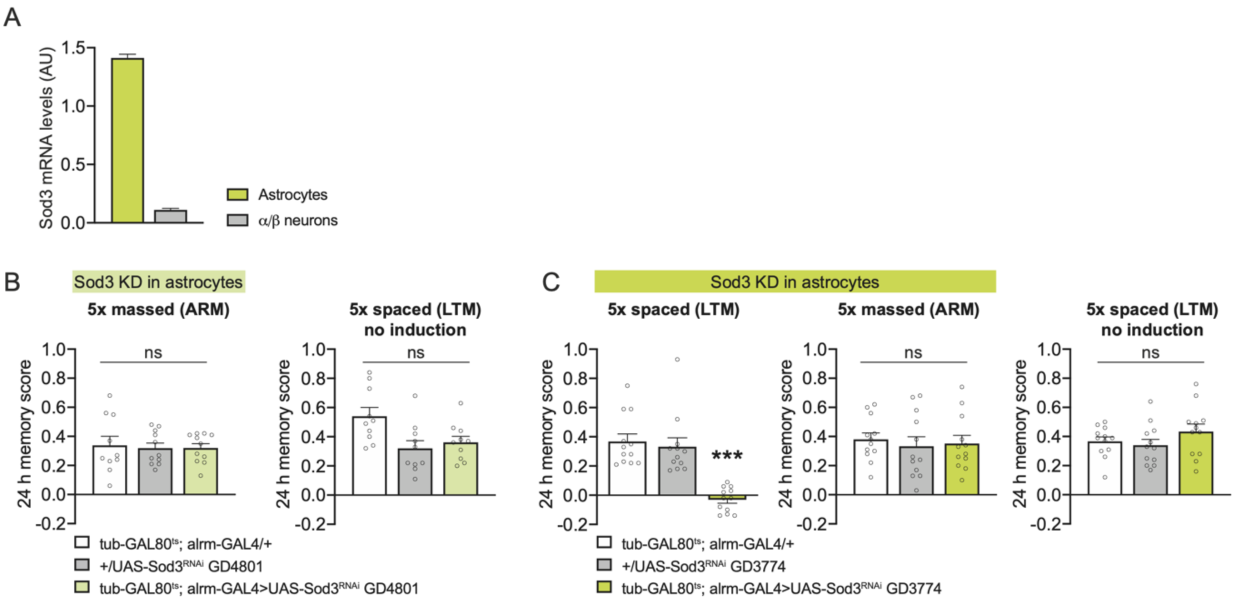


**Extended Data Fig. 2. Controls for Sod3 RNAi behavioral experiments in Fig. 1**

(**A**) Relative expression of Sod3 in astrocytes and α/β MB neurons according to published single-cell transcriptomics (*21*). Levels of Sod3 mRNA are higher in astrocytes as compared to α/β MB neurons. (**B**) Memory after massed conditioning was not affected by Sod3 KD in adult astrocytes (n = 10-11, $F_{2,29} = 0.06$, P = 0.94). Without Sod3 RNAi induction, memory after spaced training was normal (n = 10, $F_{2,27} = 5.15$, P = 0.01). (**C**) The behavioral experiments were performed with a second RNAi against Sod3. Sod3 KD in adult astrocytes impaired memory after spaced training (n = 12, $F_{2,33} = 20.88$, P < 0.001), but did not affect memory after massed training (n = 12, $F_{2,33} = 0.19$, P = 0.83). Without Sod3 RNAi induction, memory after spaced training was normal (n = 12, $F_{2,33} = 1.35$, P = 0.27).

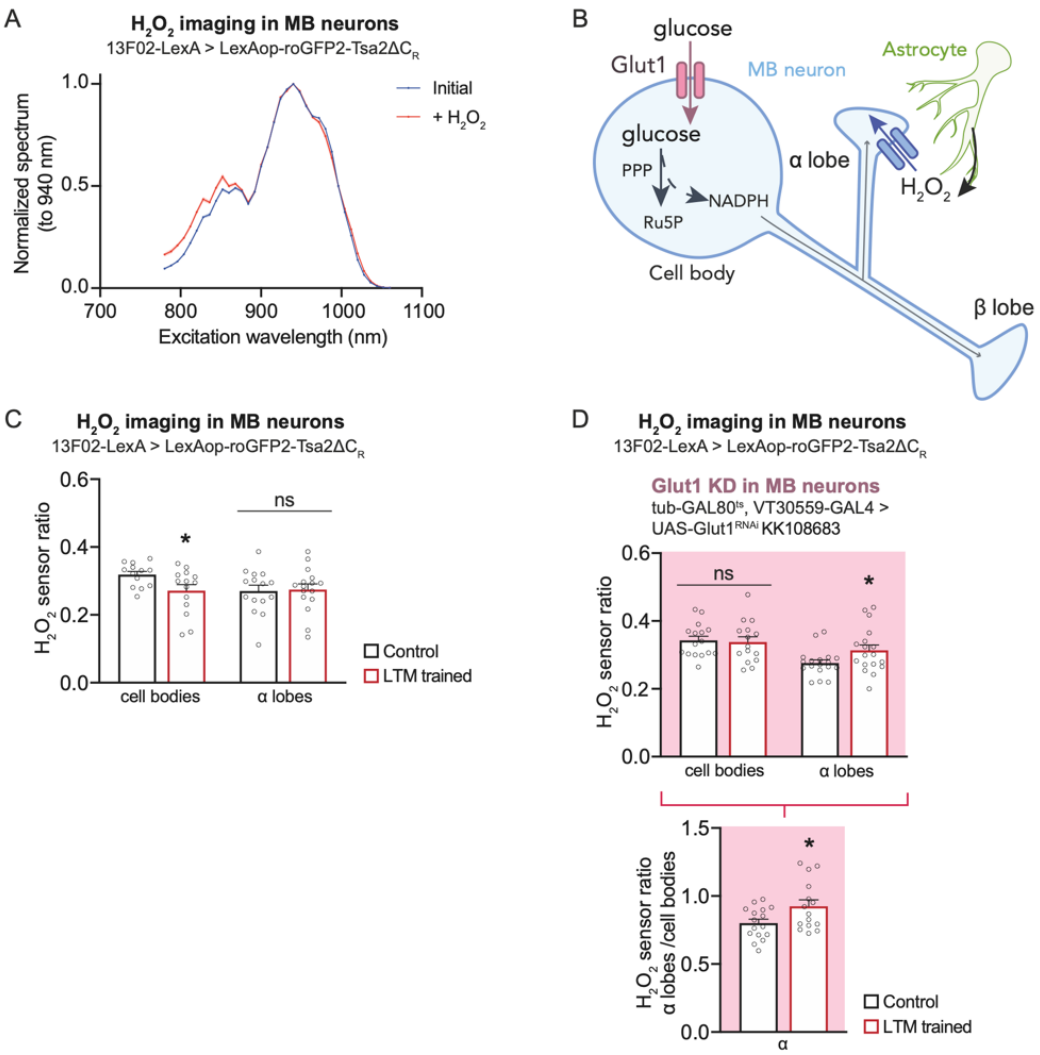


**Extended Data Fig. 3. roGFP2-Tsa2ΔC$_R$ imaging in MB neurons, related to Fig. 2.**

(**A**) Fluorescence excitation spectra of roGFP2-Tsa2ΔC$_R$ upon 2-photon excitation in basal and oxidized states. $H_2O_2$-induced oxidation of the probe led to an increase in the excitation spectrum at 780 nm and a slight decrease at 988 nm, allowing ratiometric measurement of the oxidation level of the probe (ratio 780 nm/988 nm). (**B**) Intracellular pathways recruited during LTM formation in MB neurons. Upon spaced training, glucose intake in MB neuron cell bodies via the Glut1 transporter is increased to fuel the PPP, which produces reductive power in the form of NADPH (*26*). (**C**) Measurements of $H_2O_2$ levels in cell bodies and α lobes using the roGFP2-Tsa2ΔC$_R$ $H_2O_2$ sensor expressed in MB neurons. $H_2O_2$ level (measured as

780/988 nm ratio) decreased in cell bodies upon spaced training (n = 13-14, $t_{25}$ = 2.27, P = 0.03), while it did not vary in α lobes (n = 15-16, $t_{29}$ = 0.18, P = 0.86). (**D**) Glut1 KD in MB neurons abolished the decrease in $H_2O_2$ level in cell bodies (n = 15-16, $t_{29}$ = 0.25, P = 0.81, top panel), revealing an increase in $H_2O_2$ level in α lobes after spaced training (n = 18, $t_{34}$ = 2.06, P = 0.047, top panel). The $H_2O_2$ level in α lobes normalized to the cell bodies value is increased upon spaced training when Glut1 was knocked down in adult MB neurons (n = 15-16, $t_{29}$ = 2.66, P = 0.013, bottom panel).

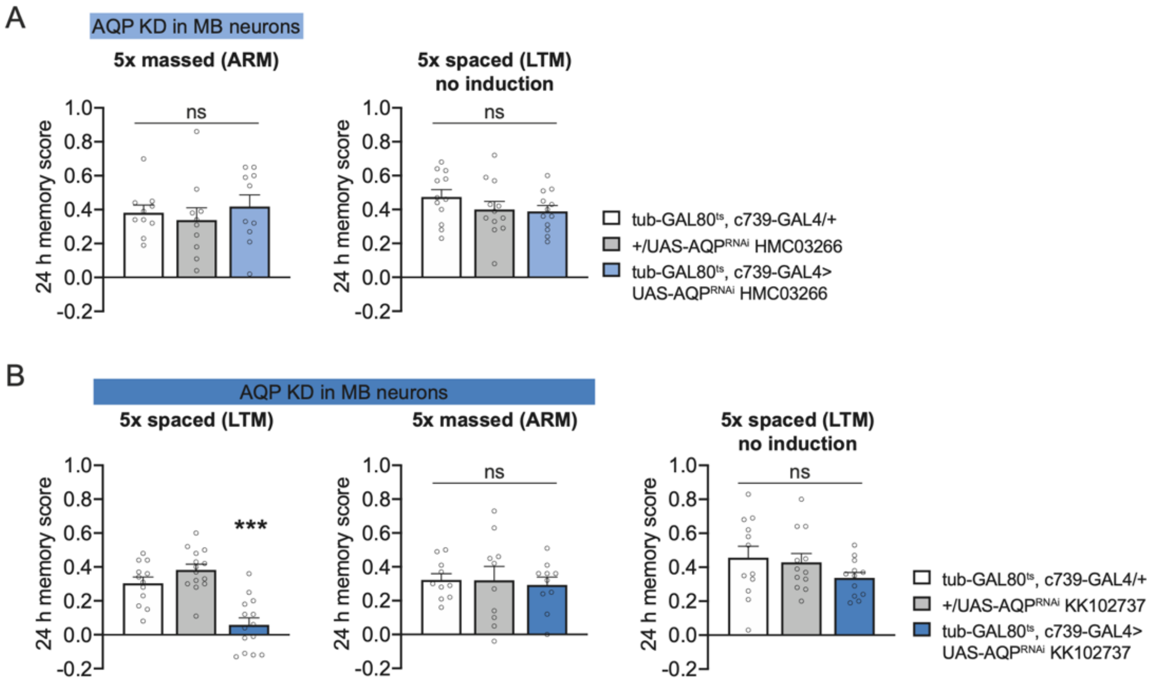


**Extended Data Fig. 4. Controls for AQP RNAi behavioral experiments in Fig. 3.**

(**A**) Memory after massed conditioning was not affected by AQP KD in adult α/β MB neurons (n = 10, $F_{2,27}$ = 0.40, P = 0.68). Without AQP RNAi induction, memory after spaced training was normal (n = 12, $F_{2,33}$ = 1.20, P = 0.32). (**B**) The behavioral experiments were performed with a second RNAi against AQP. AQP KD in adult α/β MB neurons impaired memory after spaced training (n = 12-14, $F_{2,37}$ = 21.55, P < 0.001), but did not affect memory after massed training (n = 10, $F_{2,27}$ = 0.08, P = 0.92). Without AQP RNAi induction, memory after spaced training was normal (n = 12, $F_{2,33}$ = 1.40, P = 0.26).

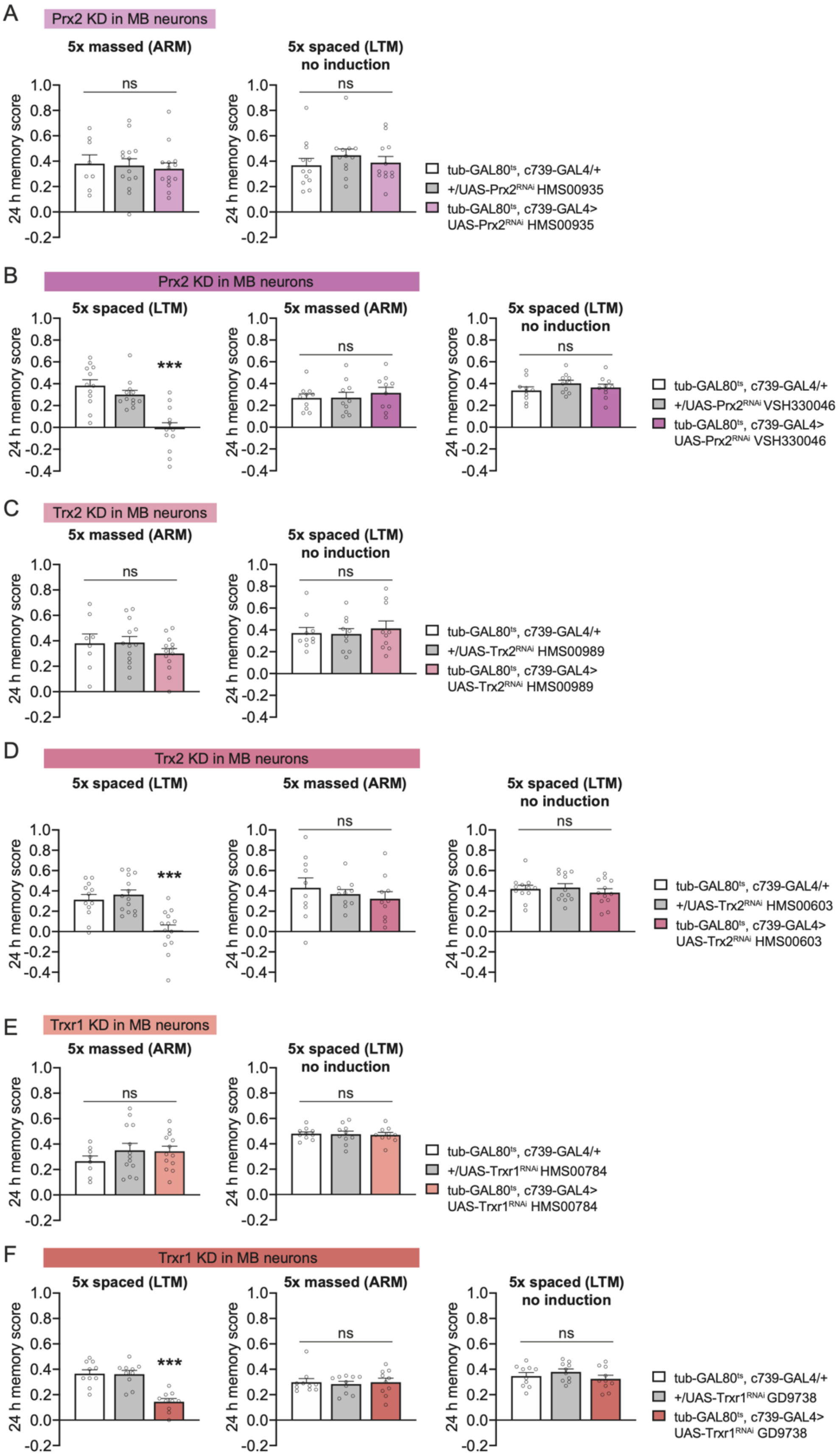
A
Prx2 KD in MB neurons
5x massed (ARM)
5x spaced (LTM) no induction
24 h memory score
ns
tub-GAL80ts, c739-GAL4/+
+/UAS-Prx2RNAi HMS00935
tub-GAL80ts, c739-GAL4> UAS-Prx2RNAi HMS00935
B
Prx2 KD in MB neurons
5x spaced (LTM)
5x massed (ARM)
5x spaced (LTM) no induction
***
tub-GAL80ts, c739-GAL4/+
+/UAS-Prx2RNAi VSH330046
tub-GAL80ts, c739-GAL4> UAS-Prx2RNAi VSH330046
C
Trx2 KD in MB neurons
5x massed (ARM)
5x spaced (LTM) no induction
tub-GAL80ts, c739-GAL4/+
+/UAS-Trx2RNAi HMS00989
tub-GAL80ts, c739-GAL4> UAS-Trx2RNAi HMS00989
D
Trx2 KD in MB neurons
5x spaced (LTM)
5x massed (ARM)
5x spaced (LTM) no induction
tub-GAL80ts, c739-GAL4/+
+/UAS-Trx2RNAi HMS00603
tub-GAL80ts, c739-GAL4> UAS-Trx2RNAi HMS00603
E
Trxr1 KD in MB neurons
5x massed (ARM)
5x spaced (LTM) no induction
tub-GAL80ts, c739-GAL4/+
+/UAS-Trxr1RNAi HMS00784
tub-GAL80ts, c739-GAL4> UAS-Trxr1RNAi HMS00784
F
Trxr1 KD in MB neurons
5x spaced (LTM)
5x massed (ARM)
5x spaced (LTM) no induction
tub-GAL80ts, c739-GAL4/+
+/UAS-Trxr1RNAi GD9738
tub-GAL80ts, c739-GAL4> UAS-Trxr1RNAi GD9738

**Extended Data Fig. 5. Controls for behavioral experiments in Fig. 3.**

(**A**) Memory after massed conditioning was not affected by Prx2 KD in adult α/β MB neurons (n = 8-14, $F_{2,33}$ = 0.13, P = 0.88). Without Prx2 RNAi induction, memory after spaced training was normal (n = 12, $F_{2,33}$ = 0.64, P = 0.53). (**B**) The behavioral experiments were performed with a second RNAi against Prx2. Prx2 KD in adult α/β MB neurons impaired memory after spaced training (n = 12, $F_{2,33}$ = 16.48, P < 0.001), but did not affect memory after massed training (n = 10, $F_{2,27}$ = 0.33, P = 0.73). Without Prx2 RNAi induction, memory after spaced training was normal (n = 10, $F_{2,27}$ = 1.09, P = 0.35). (**C**) Memory after massed conditioning was not affected by Trx2 KD in adult α/β MB neurons (n = 8-13, $F_{2,31}$ = 0.99, P = 0.38). Without Trx2 RNAi induction, memory after spaced training was normal (n = 10, $F_{2,27}$ = 0.23, P = 0.80). (**D**) The behavioral experiments were performed with a second RNAi against Trx2. Trx2 KD in adult α/β MB neurons impaired memory after spaced training (n = 12-14, $F_{2,37}$ = 14.43, P < 0.001), but did not affect memory after massed training (n = 10, $F_{2,27}$ = 0.54, P = 0.59). Without Trx2 RNAi induction, memory after spaced training was normal (n = 12, $F_{2,33}$ = 0.47, P = 0.63). (**E**) Memory after massed conditioning was not affected by Trxr1 KD in adult α/β MB neurons (n = 8-13, $F_{2,31}$ = 0.81, P = 0.45). Without Trxr1 RNAi induction, memory after spaced training was normal (n = 10, $F_{2,27}$ = 0.05, P = 0.95). (**F**) The behavioral experiments were performed with a second RNAi against Trxr1. Trxr1 KD in adult α/β MB neurons impaired memory after spaced training (n = 10, $F_{2,27}$ = 20.30, P < 0.001), but did not affect memory after massed training (n = 10, $F_{2,27}$ = 0.09, P = 0.92). Without Trxr1 RNAi induction, memory after spaced training was normal (n = 10, $F_{2,27}$ = 1.07, P = 0.36).

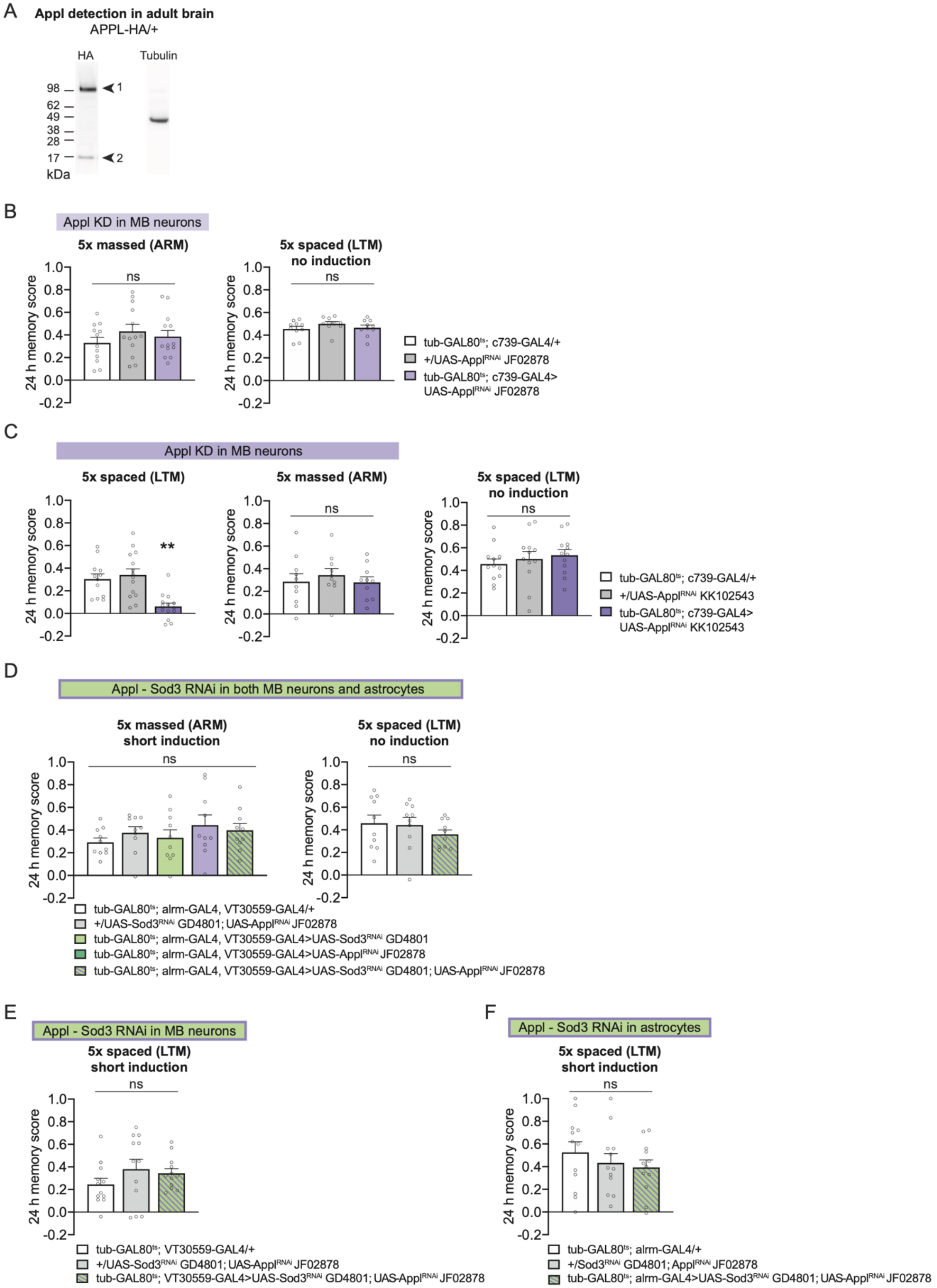


**Extended Data Fig. 6. Controls for behavioral experiments in Fig. 4.**

(**A**) Western blot against HA-tagged Appl. The expected size bands are observed at 99 kDa (label 1) and 15 kDa (label 2), corresponding to full-length APPL and the CTF-α fragment, respectively. CTF-α is generated by α-secretase cleavage of full-length APPL (*56*). Tubulin was used as a loading control. (**B**) Memory after massed conditioning was not affected by Appl KD in adult α/β MB neurons (n = 12-13, $F_{2,35}$ = 0.84, P = 0.44). Without Appl RNAi induction, memory after spaced training was normal (n = 10, $F_{2,27}$ = 1.19, P = 0.32). (**C**) The behavioral experiments were performed with a second RNAi against Appl. Appl KD in adult α/β MB neurons impaired memory after spaced training (n = 12-14, $F_{2,37}$ = 11.84, P < 0.001), but did not affect memory after massed training (n = 10, $F_{2,27}$ = 0.36, P = 0.70). Without APPL RNAi induction, memory after spaced training was normal (n = 12, $F_{2,33}$ = 0.52, P = 0.60). (**D**) Mild decreases in both Appl and Sod3 expression in adult MB neurons and astrocytes did not affect memory after massed training (n = 10, $F_{4,45}$ = 0.83, P = 0.515). Without Appl and Sod3 RNAi induction, memory after spaced training was normal (n = 10, $F_{2,27}$ = 0.72, P = 0.50). (**E**) Mild decreases in both Appl and Sod3 expression in adult MB neurons alone did not affect memory after spaced training (n = 12, $F_{2,33}$ = 1.18, P = 0.32). (**F**) Mild decreases in both Appl and Sod3 expression in adult astrocytes alone did not affect memory after spaced training (n = 12, $F_{2,33}$ = 0.70, P = 0.50).

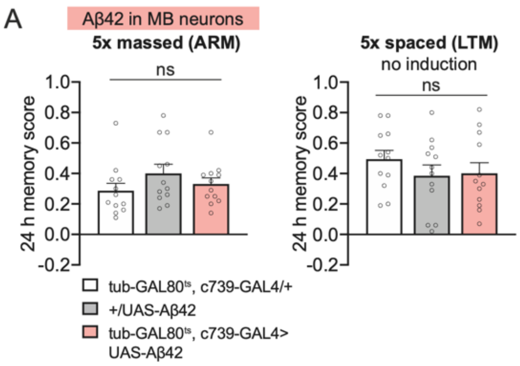


**Extended Data Fig. 7. Controls for behavioral experiments in Fig. 5.**

(**A**) Memory after massed conditioning was not affected by Aß42 expression in adult MB neurons (n=12, $F_{2,33}$=1.31, P=0.28). Without Aß42 induction, memory after spaced training was normal (n= 12, $F_{2,33}$= 0.77, P=0.47).

**Extended Data Table 1. Sensory acuity controls related to Fig. 1 and Extended Data fig. 1 and 2.**

| Genotypes | Shock reactivity | | Olfactory acuity | | | |
|---|---|---|---|---|---|---|
| | | | Octanol | | Methylcyclohexanol | |
| | Mean ± SEM | Statistics | Mean ± SEM | Statistics | Mean ± SEM | Statistics |
| tub-GAL80$^{ts}$; alrm-GAL4/+ | 0.43 ± 0.058 | n = 8<br>$F_{2,21}$ = 2.46<br>P = 0.11 | 0.59 ± 0.049 | n = 10<br>$F_{2,27}$ = 0.15<br>P = 0.87 | 0.68 ± 0.062 | n = 10<br>$F_{2,27}$ = 1.07<br>P = 0.36 |
| +/UAS-Nox RNAi HMS00429 | 0.43 ± 0.066 | | 0.60 ± 0.069 | | 0.78 ± 0.041 | |
| tub-GAL80$^{ts}$; alrm-GAL4> UAS- Nox RNAi HMS00429 | 0.61 ± 0.070 | | 0.64 ± 0.082 | | 0.67 ± 0.072 | |
| tub-GAL80$^{ts}$; alrm-GAL4/+ | 0.49 ± 0.062 | n = 12<br>$F_{2,33}$ = 0.72<br>P = 0.49 | 0.45 ± 0.029 | n = 12<br>$F_{2,33}$ = 0.015<br>P = 0.98 | 0.44 ± 0.021 | n =12<br>$F_{2,33}$ = 0.030<br>P = 0.97 |
| +/UAS-Nox RNAi KK111991 | 0.44 ± 0.045 | | 0.45 ± 0.028 | | 0.44 ± 0.038 | |
| tub-GAL80$^{ts}$; alrm-GAL4> UAS-Nox RNAi KK111991 | 0.53 ± 0.064 | | 0.46 ± 0.026 | | 0.45 ± 0.029 | |
| tub-GAL80$^{ts}$; alrm-GAL4/+ | 0.39 ± 0.053 | n = 8<br>$F_{2,21}$ = 1.98 | 0.74 ± 0.042 | n = 10<br>$F_{2,27}$ = 0.20 | 0.77 ± 0.066 | n = 10<br>$F_{2,27}$ = 0.76 |

| | | | | | | |
|---|---|---|---|---|---|---|
| +/UAS-G6PD RNAi KK108898 | 0.31 ± 0.056 | P = 0.16 | 0.72 ± 0.053 | P = 0.82 | 0.68 ± 0.043 | P = 0.48 |
| tub-GAL80$^{ts}$; alrm-GAL4> UAS-G6PD RNAi KK108898 | 0.48 ± 0.072 | | 0.68 ± 0.082 | | 0.68 ± 0.070 | |
| tub-GAL80$^{ts}$; alrm-GAL4/+ | 0.40 ± 0.066 | n = 8 $F_{2,21}$ = 1.62 P = 0.22 | 0.61 ± 0.074 | n = 10 $F_{2,27}$ = 0.87 P = 0.43 | 0.77 ± 0.080 | n = 10 $F_{2,27}$ = 0.11 P = 0.89 |
| +/UAS-Pgd RNAi HMC05959 | 0.31 ± 0.064 | | 0.73 ± 0.062 | | 0.77 ± 0.039 | |
| tub-GAL80$^{ts}$; alrm-GAL4> UAS-Pgd RNAi HMC05959 | 0.46 ± 0.047 | | 0.65 ± 0.065 | | 0.73 ± 0.092 | |
| tub-GAL80$^{ts}$; alrm-GAL4/+ | 0.63 ± 0.044 | n = 12 $F_{2,33}$ = 1.01 P = 0.37 | 0.72 ± 0.034 | n = 12 $F_{2,33}$ = 0.006 P >0.99 | 0.79 ± 0.038 | n = 12 $F_{2,33}$ = 0.155 P = 0.86 |
| +/UAS-Pgls RNAi HMS02626 | 0.59 ± 0.052 | | 0.73 ± 0.035 | | 0.78 ± 0.028 | |
| tub-GAL80$^{ts}$; alrm-GAL4> Pgls RNAi HMS02626 | 0.69 ± 0.045 | | 0.72 ± 0.032 | | 0.81 ± 0.024 | |

| | | | | | | |
|---|---|---|---|---|---|---|
| tub-GAL80$^{ts}$; alrm-GAL4/+ | 0.68 ± 0.047 | n = 10 $F_{2,27} = 0.77$ P = 0.47 | 0.67 ± 0.036 | n = 10 $F_{2,27} = 1.64$ P = 0.21 | 0.80 ± 0.062 | n = 10 $F_{2,27} = 3.16$ P = 0.06 |
| +/UAS-nAChRα7 RNAi JF02570 | 0.57 ± 0.069 | | 0.60 ± 0.037 | | 0.61 ± 0.050 | |
| tub-GAL80$^{ts}$; alrm-GAL4> UAS-nAChRα7 RNAi JF02570 | 0.62 ± 0.065 | | 0.58 ± 0.040 | | 0.64 ± 0.058 | |
| tub-GAL80$^{ts}$; alrm-GAL4/+ | 0.41 ± 0.049 | n = 12 $F_{2,33} = 0.29$ P = 0.75 | 0.50 ± 0.019 | n = 12 $F_{2,33} = 0.51$ P = 0.60 | 0.54 ± 0.050 | n = 12 $F_{2,33} = 0.18$ P = 0.84 |
| +/UAS-nAChRα7 RNAi KK108471 | 0.45 ± 0.054 | | 0.54 ± 0.030 | | 0.51 ± 0.033 | |
| tub-GAL80$^{ts}$; alrm-GAL4> UAS-nAChRα7 RNAi KK108471 | 0.46 ± 0.053 | | 0.51 ± 0.047 | | 0.50 ± 0.047 | |
| tub-GAL80$^{ts}$; alrm-GAL4/+ | 0.43 ± 0.083 | n = 8 $F_{2,21} = 0.30$ P = 0.75 | 0.67 ± 0.068 | n = 10 $F_{2,27} = 0.20$ P = 0.82 | 0.78 ± 0.066 | n = 10 $F_{2,27} = 0.63$ P = 0.54 |
| +/UAS-Sod3 RNAi GD4801 | 0.44 ± 0.062 | | 0.67 ± 0.056 | | 0.67 ± 0.073 | |
| tub-GAL80$^{ts}$; alrm-GAL4> | 0.50 ± 0.064 | | 0.62 ± 0.064 | | 0.75 ± 0.067 | |

| | | | | | | |
|---|---|---|---|---|---|---|
| UAS-Sod3 RNAi GD4801 | | | | | | |
| tub-GAL80$^{ts}$; alrm-GAL4/+ | 0.59 ± 0.054 | | 0.66 ± 0.028 | | 0.80 ± 0.037 | |
| +/UAS-Sod3 RNAi GD3774 | 0.58 ± 0.074 | n = 12<br>$F_{2,33}$ = 0.41<br>P = 0.67 | 0.74 ± 0.033 | n = 12<br>$F_{2,33}$ = 1.60<br>P = 0.22 | 0.83 ± 0.023 | n = 12<br>$F_{2,33}$ = 0.77<br>P = 0.47 |
| tub-GAL80$^{ts}$; alrm-GAL4> UAS-Sod3 RNAi GD3774 | 0.65 ± 0.063 | | 0.69 ± 0.030 | | 0.77 ± 0.039 | |

**Extended Dara Table 2. Sensory acuity controls related to Fig. 3 and Extended Data fig. 4.**

| Genotypes | Shock reactivity | | Olfactory acuity | | | |
|---|---|---|---|---|---|---|
| | | | Octanol | | Methylcyclohexanol | |
| | Mean ± SEM | Statistics | Mean ± SEM | Statistics | Mean ± SEM | Statistics |
| tub-GAL80$^{ts}$; c739-GAL4/+ | 0.70 ± 0.057 | n = 12<br>$F_{2,33}$ = 2.49<br>P = 0.10 | 0.49 ± 0.047 | n = 10<br>$F_{2,27}$ = 0.31<br>P = 0.73 | 0.50 ± 0.061 | n = 10<br>$F_{2,27}$ = 0.06<br>P = 0.94 |
| +/UAS-AQP RNAi HMC03266 | 0.55 ± 0.079 | | 0.47 ± 0.050 | | 0.52 ± 0.069 | |
| tub-GAL80$^{ts}$; c739-GAL4> UAS-AQP RNAi HMC03266 | 0.75 ± 0.062 | | 0.52 ± 0.052 | | 0.53 ± 0.058 | |
| tub-GAL80$^{ts}$; c739-GAL4/+ | 0.66 ± 0.055 | n = 12<br>$F_{2,33}$ = 0.17<br>P = 0.85 | 0.60 ± 0.054 | n = 10<br>$F_{2,27}$ = 0.41<br>P = 0.67 | 0.69 ± 0.060 | n = 10<br>$F_{2,27}$ = 0.48<br>P = 0.62 |
| +/UAS-AQP RNAi KK102737 | 0.68 ± 0.071 | | 0.55 ± 0.054 | | 0.63 ± 0.037 | |
| tub-GAL80$^{ts}$; c739-GAL4> UAS-AQP RNAi KK102737 | 0.71 ± 0.060 | | 0.53 ± 0.043 | | 0.67 ± 0.032 | |

**Extended Data Table 3. Sensory acuity controls related to Fig. 3 and Extended Data fig. 5.**

| Genotypes | Shock reactivity | | Olfactory acuity | | | |
|---|---|---|---|---|---|---|
| | | | Octanol | | Methylcyclohexanol | |
| | Mean ± SEM | Statistics | Mean ± SEM | Statistics | Mean ± SEM | Statistics |
| tub-GAL80$^{ts}$; c739-GAL4/+ | 0.57 ± 0.045 | n = 12<br>$F_{2,33} = 1.91$<br>P = 0.16 | 0.67 ± 0.058 | n = 10<br>$F_{2,27} = 0.42$<br>P = 0.66 | 0.63 ± 0.054 | n = 10<br>$F_{2,27} = 2.71$<br>P = 0.08 |
| +/UAS-Prx2 RNAi HMS00935 | 0.69 ± 0.049 | | 0.64 ± 0.040 | | 0.78 ± 0.039 | |
| tub-GAL80$^{ts}$; c739-GAL4> UAS-Prx2 RNAi HMS00935 | 0.63 ± 0.035 | | 0.61 ± 0.044 | | 0.70 ± 0.049 | |
| tub-GAL80$^{ts}$; c739-GAL4/+ | 0.57 ± 0.044 | n = 12<br>$F_{2,33} = 0.42$<br>P = 0.66 | 0.52 ± 0.027 | n = 10<br>$F_{2,27} = 0.52$<br>P = 0.60 | 0.47 ± 0.029 | n = 10<br>$F_{2,27} = 1.60$<br>P = 0.22 |
| +/UAS-Prx2 RNAi VSH330046 | 0.56 ± 0.028 | | 0.54 ± 0.021 | | 0.49 ± 0.020 | |
| tub-GAL80$^{ts}$; c739-GAL4> UAS-Prx2 RNAi VSH330046 | 0.61 ± 0.039 | | 0.50 ± 0.022 | | 0.54 ± 0.026 | |
| tub-GAL80$^{ts}$; c739-GAL4/+ | 0.45 ± 0.059 | n = 12<br>$F_{2,33} = 0.82$ | 0.55 ± 0.041 | n = 12<br>$F_{2,33} = 3.43$ | 0.68 ± 0.060 | n = 12<br>$F_{2,33} = 1.22$ |

| | | | | | | |
|---|---|---|---|---|---|---|
| +/UAS-Trx2 RNAi HMS00989 | 0.54 ± 0.077 | P = 0.45 | 0.67 ± 0.040 | P = 0.04 | 0.71 ± 0.041 | P = 0.31 |
| tub-GAL80$^{ts}$; c739-GAL4> UAS-Trx2 RNAi HMS00989 | 0.55 ± 0.048 | | 0.56 ± 0.029 | | 0.60 ± 0.050 | |
| tub-GAL80$^{ts}$; c739-GAL4/+ | 0.58 ± 0.039 | n = 12<br>$F_{2,33}$ = 0.21<br>P = 0.81 | 0.64 ± 0.046 | n = 12<br>$F_{2,33}$ = 0.11<br>P = 0.89 | 0.66 ± 0.062 | n = 12<br>$F_{2,33}$ = 1.12<br>P = 0.34 |
| +/UAS-Trx2 RNAi HMS00603 | 0.62 ± 0.065 | | 0.64 ± 0.047 | | 0.75 ± 0.050 | |
| tub-GAL80$^{ts}$; c739-GAL4> UAS-Trx2 RNAi HMS00603 | 0.58 ± 0.070 | | 0.62 ± 0.035 | | 0.64 ± 0.044 | |
| tub-GAL80$^{ts}$; c739-GAL4/+ | 0.48 ± 0.077 | n = 10<br>$F_{2,27}$ = 0.52<br>P = 0.60s | 0.71 ± 0.044 | n = 10<br>$F_{2,27}$ = 0.90<br>P = 0.42 | 0.70 ± 0.038 | n = 10<br>$F_{2,27}$ = 1.47<br>P = 0.25 |
| +/UAS-Trxr1 RNAi HMS00784 | 0.56 ± 0.059 | | 0.69 ± 0.036 | | 0.75 ± 0.037 | |
| tub-GAL80$^{ts}$; c739-GAL4> UAS-Trxr1 RNAi HMS00784 | 0.47 ± 0.077 | | 0.78 ± 0.062 | | 0.65 ± 0.042 | |

| | | | | | | |
|---|---|---|---|---|---|---|
| tub-GAL80$^{ts}$; c739-GAL4/+ | 0.49 ± 0.049 | n = 12<br>$F_{2,33}$ = 0.13<br>P = 0.88 | 0.67 ± 0.060 | n = 10<br>$F_{2,27}$ = 0.08<br>P = 0.92 | 0.60 ± 0.036 | n = 10<br>$F_{2,27}$ = 0.34<br>P = 0.71 |
| +/UAS-Trxr1 RNAi GD9738 | 0.52 ± 0.041 | | 0.66 ± 0.058 | | 0.67 ± 0.060 | |
| tub-GAL80$^{ts}$; c739-GAL4> UAS-Trxr1 RNAi GD9738 | 0.50 ± 0.048 | | 0.64 ± 0.049 | | 0.65 ± 0.062 | |

**Extended Data Table 4. Sensory acuity controls related to Fig. 4 and Extended Data fig. 6.**

| Genotypes | Shock reactivity | | Olfactory acuity | | | |
|---|---|---|---|---|---|---|
| | | | Octanol | | Methylcyclohexanol | |
| | Mean ± SEM | Statistics | Mean ± SEM | Statistics | Mean ± SEM | Statistics |
| tub-GAL80$^{ts}$; c739-GAL4/+ | 0.53 ± 0.055 | n = 14<br>$F_{2,39} = 1.36$<br>P = 0.27 | 0.74 ± | n = 10<br>$F_{2,27} = 0.12$<br>P = 0.88 | 0.64 ± | n = 10<br>$F_{2,27} = 3.84$<br>P = 0.03 |
| +/UAS-Appl RNAi JF02878 | 0.64 ± 0.048 | | 0.73 ± | | 0.73 ± | |
| tub-GAL80$^{ts}$; c739-GAL4> UAS- Appl RNAi JF02878 | 0.62 ± 0.041 | | 0.71 ± | | 0.84 ± | |
| tub-GAL80$^{ts}$; c739-GAL4/+ | 0.64 ± 0.077 | n = 12<br>$F_{2,33} = 0.61$<br>P = 0.55 | 0.60 ± 0.053 | n = 10<br>$F_{2,27} = 1.19$<br>P = 0.32 | 0.82 ± 0.034 | n = 10<br>$F_{2,27} = 0.49$<br>P = 0.62 |
| +/UAS-Appl RNAi KK102543 | 0.53 ± 0.073 | | 0.58 ± 0.067 | | 0.79 ± 0.032 | |
| tub-GAL80$^{ts}$; c739-GAL4> UAS- Appl RNAi KK102543 | 0.57 ± 0.065 | | 0.49 ±0.037 | | 0.76 ± 0.55 | |
| tub-GAL80$^{ts}$; alrm-GAL4, VT30559-GAL4/+ | 0.59 ± 0.031 | n = 10<br>$F_{2,27} = 0.75$<br>P = 0.48 | 0.52 ± 0.055 | n = 10<br>$F_{2,27} = 3.14$<br>P = 0.06 | 0.81 ± 0.052 | n = 10<br>$F_{2,27} = 1.03$<br>P =0.37 |
| +/ UAS-Sod3 RNAi GD4801; UAS-Appl RNAi KK102543 | 0.58 ± 0.049 | | 0.66 ± 0.047 | | 0.84 ± 0.033 | |
| tub-GAL80$^{ts}$; alrm-GAL4, VT30559-GAL4> UAS-Sod3 RNAi GD4801; UAS-Appl RNAi KK102543 | 0.66 ± 0.073 | | 0.51 ± 0.040 | | 0.76 ± 0.034 | |

**Extended data Table 5. Sensory acuity controls related to Fig. 5 and Extended Data fig. 7.**

| Genotypes | Shock reactivity | | Olfactory acuity | | | |
|---|---|---|---|---|---|---|
| | | | Octanol | | Methylcyclohexanol | |
| | Mean ± SEM | Statistics | Mean ± SEM | Statistics | Mean ± SEM | Statistics |
| tub-GAL80$^{ts}$; c739-GAL4/+ | 0.55 ± 0.081 | n = 10<br>$F_{2,27}$ = 0.95<br>P = 0.40 | 0.72 ± 0.047 | n = 12<br>$F_{2,33}$ = 2.30<br>P = 0.12 | 0.72 ± 0.052 | n = 12<br>$F_{2,33}$ = 1.63<br>P = 0.21 |
| +/UAS-Aß42 | 0.41 ± 0.054 | | 0.69 ± 0.048 | | 0.59 ± 0.052 | |
| tub-GAL80$^{ts}$; c739-GAL4> UAS-Aß42 | 0.48 ± 0.067 | | 0.57 ± 0.059 | | 0.62 ± 0.061 | |
