## Supplementary Table for "Long-term memory formation depends on an astrocyte-to-neuron H_2_O_2_ signaling"

**Supplementary Table 1:** ***Drosophila melanogaster*** **strains used in the study**

| **Strain** | **Source / reference** |
|---|---|
| tub-GAL80$^{ts}$; alrm-GAL4 | de Tredern et al. 2021 |
| tub-GAL80$^{ts}$; c739-GAL4 | Turrel et al. 2018 |
| tub-GAL80$^{ts}$, 13F02-LexA; VT30559-GAL4 | de Tredern et al. 2021 |
| tub-GAL80$^{ts}$, 13F02-LexA; alrm-GAL4 | This study |
| tub-GAL80$^{ts}$; alrm-GAL4, VT30559-GAL4 | This study |
| 86E01-LexA | Bloomington Drosophila Stock Center (BDSC) 54287 |
| tub-GAL80$^{ts}$, 86E01-LexA; VT30559-GAL4 | This study |
| UAS-Nox$^{RNAi}$ HMS00429 | BDSC 32433 |
| UAS-Nox$^{RNAi}$ KK111991 | Vienna Drosophila Resource Center (VDRC) 102559 |
| UAS-G6PD$^{RNAi}$ KK108898 | VDRC 101507 |
| UAS-Pgd$^{RNAi}$ HMC05959 | BDSC 65078 |
| UAS-Pgls$^{RNAi}$ HMS02626 | BDSC 42933 |
| UAS-nAChRa7$^{RNAi}$ JF02570 | BDSC 27251 |
| UAS-nAChRa7$^{RNAi}$ KK108471 | VDRC 100756 |
| UAS-Sod3$^{RNAi}$ GD4801 | VDRC 37793 |
| UAS-Sod3$^{RNAi}$ GD3774 | VDRC 8760 |
| UAS-Glut1$^{RNAi}$ KK108683 | VDRC 101365 |
| UAS-AQP$^{RNAi}$ HMC03266 | BDSC 51504 |
| UAS-AQP$^{RNAi}$ KK102737 | VDRC 109314 |
| UAS-Prx2$^{RNAi}$ HMS00935 | BDSC 34971 |
| UAS-Prx2$^{RNAi}$ VSH330046 | VDRC 330046 |
| UAS-Trx2$^{RNAi}$ HMS00989 | BDSC 34019 |
| UAS-Trx2$^{RNAi}$ HMS00603 | BDSC 33721 |
| UAS-Trxr1$^{RNAi}$ HMS00784 | BDSC 32984 |
| UAS-Trxr1$^{RNAi}$ GD9738 | VDRC 47308 |
| UAS-Appl$^{RNAi}$ JF02878 | BDSC 28043 |

| | |
|---|---|
| UAS-Appl$^{RNAi}$ KK102543 | VDRC 108312 |
| UAS-Aß42 | Finelli et al. 2004 |
| UAS-GCaMP6f | BDSC 42747 |
| LexAop-GCaMP6f | BDSC 44277 |
| LexAop-GCaMP6f; UAS-Aß42 | This study |
| UAS-GCaMP6f; UAS-nAChRa7$^{RNAi}$ JF02570 | This study |
| LexAop-roGFP-Tsa2DC$_R$ | This study |
| LexAop-roGFP-Tsa2DC$_R$; UAS-Glut1$^{RNAi}$ KK108683 | This study |
| LexAop-roGFP-Tsa2DC$_R$; UAS-Nox$^{RNAi}$ HMS00429 | This study |
| LexAop-roGFP-Tsa2DC$_R$; UAS-G6PD$^{RNAi}$ KK108898 | This study |
| LexAop-roGFP-Tsa2DC$_R$; UAS-nAChRa7$^{RNAi}$ JF02570 | This study |
| LexAop-roGFP-Tsa2DC$_R$; UAS-Sod3$^{RNAi}$ GD4801 | This study |
| LexAop-roGFP-Tsa2DC$_R$; UAS-AQP$^{RNAi}$ HMC03266 | This study |
| LexAop-roGFP-Tsa2DC$_R$; UAS-Appl$^{RNAi}$ JF02878 | This study |
| LexAop-roGFP-Tsa2DC$_R$; UAS-Aß42 | This study |
| APPL-HA | This study |
